# THEM6-mediated lipid remodelling sustains stress resistance in cancer

**DOI:** 10.1101/2021.04.04.438378

**Authors:** Arnaud Blomme, Coralie Peter, Ernest Mui, Giovanny Rodriguez Blanco, Louise M. Mason, Lauren E. Jamieson, Grace H. McGregor, Sergio Lilla, Chara Ntala, Rachana Patel, Marc Thiry, Sonia H. Y. Kung, Catriona A. Ford, Linda K. Rushworth, David J. McGarry, Susan Mason, Peter Repiscak, Colin Nixon, Mark J. Salji, Elke Markert, Gillian M. MacKay, Jurre J. Kamphorst, Duncan Graham, Karen Faulds, Ladan Fazli, Martin E. Gleave, Edward Avezov, Joanne Edwards, Huabing Yin, David Sumpton, Karen Blyth, Pierre Close, Daniel J. Murphy, Sara Zanivan, Hing Y. Leung

## Abstract

Despite the clinical benefit of androgen-deprivation therapy (ADT), the majority of patients with advanced prostate cancer (PCa) ultimately develop lethal castration-resistant prostate cancer (CRPC). In this study, we identified thioesterase superfamily member 6 (THEM6) as a marker of ADT resistance in PCa. In patients, THEM6 expression correlates with progressive disease and is associated with poor survival. THEM6 deletion reduces *in vivo* tumour growth and restores castration sensitivity in orthograft models of CRPC. Mechanistically, THEM6 is located at the endoplasmic reticulum (ER) membrane and controls lipid homeostasis by regulating intracellular levels of ether lipids. Consequently, THEM6 loss in CRPC cells significantly alters ER function, reducing *de novo* sterol biosynthesis and preventing lipid-mediated induction of ATF4. Finally, we show that THEM6 is required for the establishment of the MYC-induced stress response. Thus, similar to PCa, THEM6 loss significantly impairs tumorigenesis in the MYC-dependent subtype of triple negative breast cancer. Altogether, our results highlight THEM6 as a novel component of the treatment-induced stress response and a promising target for the treatment of CRPC and MYC-driven cancer.

## INTRODUCTION

Androgen deprivation therapy (ADT) and androgen receptor (AR)-targeted therapies have significantly improved outcomes for patients suffering from advanced prostate cancer (PCa). However, treatment resistance ultimately leads to the development of lethal castration-resistant prostate cancer (CRPC), which remains a major therapeutic challenge.

Resistance to treatment is accompanied by a plethora of cellular and metabolic adaptations that allow cancer cells to cope with stress-inducing factors (1). Together with mitochondria, the endoplasmic reticulum (ER) plays a central role in the regulation of stress-signalling pathways. Indeed, the ER is critical for the establishment of a complex stress response, termed the unfolded protein response (UPR), that orchestrates the cellular adaptation to various perturbations such as impairment of protein or lipid homeostasis. Activation of the UPR relies on the coordinated action of three major branches, each of them characterized by the activity of a specific ER stress sensor: Inositol Requiring Enzyme 1 (IRE1α), PRKR-like Endoplasmic Reticulum Kinase (PERK) and Activating Transcription Factor 6 (ATF6). In cancer, correct induction of the UPR is required to support oncogenic transformation (2). Persistent activation of ER stress responses can further promote tumour progression, metastasis dissemination and resistance to therapy (3). Therefore, clinical targeting of the UPR, alone or in combination with other treatment modalities, is regarded as a promising strategy for the treatment of aggressive cancer (4–7). This strategy is particularly effective in the context of PCa, where inhibition of the IRE1a branch of the UPR significantly impairs tumour growth and activation of c-MYC signalling (8), an important feature of ADT resistance (9). Similarly, targeting of the PERK-eIF2a-ATF4 branch of the UPR effectively reduces tumour progression and metastasis dissemination in preclinical models of CRPC (10), while ATF4 signalling is essential for PCa growth and survival (11).

ER is also the primary site of lipid and cholesterol biosynthesis. Changes in lipid homeostasis, such as impaired membrane lipid saturation (12) or imbalance in the levels of phospho- and sphingolipid species (13, 14), can also contribute to ER stress. Recently we and others demonstrated the importance of lipid remodelling in tumour resistance to antiandrogen therapy (15, 16). Thus, targeting lipid-mediated ER stress could be considered as a potential therapeutic option for the treatment of CRPC.

Acyl-CoA thioesterases (ACOTs) are a class of enzymes that hydrolyse acyl-CoA molecules. In contrast with Type I enzymes, Type II ACOTs are related by structure rather than sequence. Type II ACOTs are characterised by the presence of an evolutionarily conserved domain, the “Hotdog” domain, which confers the thioesterase enzymatic activity (17). Type II ACOTs also include members of the Thioesterase Superfamily (THEM), which primarily function to deactivate fatty acyl-CoA thioesters and generate free fatty acids (FA) and CoA. THEM proteins are involved in the regulation of intracellular FA trafficking and have been shown to influence both lipogenesis and beta-oxidation depending on the physiological context (18). Interestingly, *Them1* and *Them2* knockout mice exhibit severely attenuated responses to ER stress induction (19, 20), thus supporting the role of these proteins in lipid-mediated activation of the UPR.

In this study, we identified Thioesterase Superfamily Member 6 (THEM6) as a clinically relevant protein associated with resistance to ADT. Mechanistically, we show that THEM6 maintains lipid homeostasis by controlling intracellular levels of ether lipids. Consequently, THEM6 expression is critical for ER membrane protein trafficking, sterol biosynthesis and ATF4 activation under ADT conditions. Importantly, we further observe that *THEM6* amplification is a frequent genomic alteration in cancer, and that THEM6 is functionally required to sustain MYC-induced UPR activation. Taken together, our results highlight the potential of THEM6 as a future therapeutic option for MYC-driven cancer.

## RESULTS

### THEM6 is overexpressed in CRPC and correlates with poor clinical outcome

We previously performed an in-depth proteomic analysis comparing *in vivo* hormone-naïve (HN) and castration–resistant (CRPC) orthograft models of PCa (21). From this comparison, we identified THEM6/c8orf55 as significantly upregulated in CRPC tumours (22rv1 and LNCaP AI) in comparison to HN counterparts (CWR22res and LNCaP, respectively) (Fig. 1a and Suppl. Fig. 1a). Increased THEM6 levels in CRPC were validated *in vivo* (Fig. 1b-c) and *in vitro* (Suppl. Fig. 1b). Of note, THEM6 expression was the lowest in normal prostate epithelial cells (RWPE-1) when compared with multiple PCa cell lines (Suppl. Fig. 1b). Immunohistochemistry (IHC) staining of prostate orthografts confirmed THEM6 over-expression in CRPC tumours and highlighted strong cytoplasmic and perinuclear staining in tumour epithelial cells (Fig. 1c).

**Figure 1:**
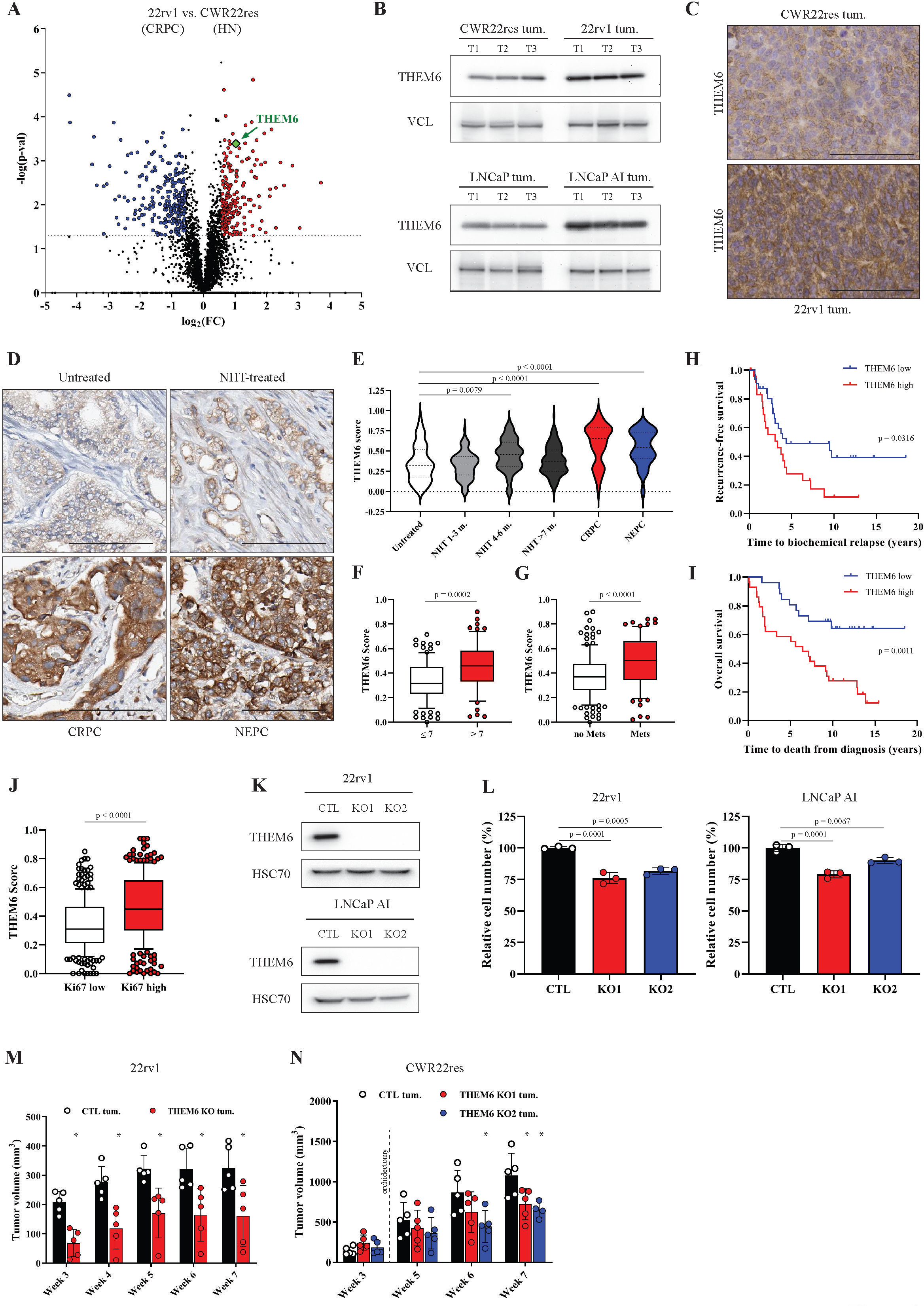
THEM6 is a clinically relevant protein overexpressed in CRPC. **A**, Volcano plot of the differentially modulated proteins in 22rv1 (CRPC) versus CWR22res (HN) tumours. Red and blue dots represent the proteins that are significantly up- and down-regulated, respectively (p-value < 0.05, FC = 1.5). **B**, Western blot analysis of THEM6 expression in CRPC (22rv1 and LNCaP AI) and HN (CWR22res and LNCaP) prostate orthografts. VCL was used as a sample loading control. **C**, IHC staining of THEM6 expression in CWR22res and 22rv1 orthografts. Scale bar represents 100 µm. **D**, IHC staining of THEM6 expression in treatment naïve, NHT-treated, CRPC and NEPC tumours. Scale bar represents 100 µm. **E**, Quantification of THEM6 staining in PCa tissue samples. **F-G**, Quantification of THEM6 expression in PCa tissue samples according to Gleason score (**F**) and metastatic status (**G**). **H-I**, Kaplan-Meier recurrence-free (**H**) and overall (**I**) survival analysis of PCa patients stratified according to median THEM6 expression. **J**, Quantification of THEM6 expression in PCa tissue samples according to Ki67 expression. **K**, Western blot analysis of THEM6 expression in CTL and THEM6 KO CRPC cells. HSC70 was used as a sample loading control. **L**, Proliferation of CTL and THEM6 KO CRPC cells after 72 hours. Data are expressed as a percentage of CTL cells. **M**, Tumour volume (measured by ultrasound) of CTL and THEM6 KO 22rv1-derived orthografts developed in surgically castrated mice. **N**, Tumour volume (measured by ultrasound) of CTL and THEM6 KO CWR22res-derived orthografts. Orchidectomy was performed 3-weeks after cell implantation. Panel **E**: Centre line corresponds to median of data, top and bottom lines correspond to upper and lower quartiles. Panels **F, G, J**: Centre line corresponds to median of data, top and bottom of box correspond to 90th and 10th percentile, respectively. Whiskers extend to adjacent values (minimum and maximum data points not considered outliers). Panels **L, M, N**: Data are presented as mean values +/- SD. <u>Statistical analysis</u> : **E**: 1-way ANOVA with a Dunnett’s multiple comparisons test. **F, G, J, M**: two-tailed Mann-Whitney U test. **H, I**: logrank test. **L**: 1-way ANOVA with a Dunnett’s multiple comparisons test. **N**: Kruskal-Wallis test.

We next assessed the clinical relevance of THEM6 in CRPC. For this purpose, we examined THEM6/c8orf55 expression in three publicly available datasets of localised and metastatic PCa (Suppl. Fig. 1c-f). Analysis of the PRAD TCGA data revealed that *THEM6* mRNA expression was significantly higher in tumour tissue than in normal prostate epithelium (Suppl. Fig. 1c). Similarly, *THEM6* levels gradually increased from benign to localised and metastatic PCa lesions ((22), Suppl. Fig. 1d), and were also elevated in distant metastases ((23), Suppl. Fig. 1e). Additionally, high *THEM6* expression was significantly associated with poor patient survival ((23), Suppl. Fig. 1f). To further demonstrate the importance of THEM6 in PCa, we performed IHC on a tissue microarray (TMA, n = 297) comprised of tumours obtained from treatment-naïve (Untreated), treatment-responsive (neoadjuvant hormonal therapy, NHT-treated), treatment-resistant (CRPC) and neuroendocrine (NEPC) PCa patients (Fig. 1d). CRPC and NEPC tumours exhibited significantly higher levels of the THEM6 protein than untreated tumours, with the highest score obtained for CRPC (Fig. 1e). High THEM6 levels significantly correlated with high Gleason score and high T-stage tumours (Fig. 1f and Suppl. Fig. 1g), and were further associated with the presence of metastases and cancer recurrence (Fig. 1g and Suppl. Fig. 1h). Finally, we validated THEM6 as a prognostic factor in PCa by staining a second TMA composed of tumour biopsies from treatment-naïve patients documented with a 20-years follow-up (n = 69). In this cohort, high THEM6 expression was strongly associated with poor patient survival (recurrence-free and overall survival, Fig. 1h-i). Finally, in both patient cohorts, high THEM6 levels were significantly associated with high Ki67 expression (Fig. 1j and Suppl. Fig. 1i), indicating that THEM6 was expressed at higher levels in highly proliferative tumours.

#### Loss of THEM6 affects CRPC tumour growth

To investigate the role of THEM6 in CRPC, we generated stable CRISPR-based THEM6 knockout cell lines (hereafter referred to as THEM6 KO) (Fig. 1k). On average, THEM6 KO resulted in a mild (∼20-25%) decrease in CRPC cell proliferation (Fig. 1l). *In vivo*, loss of THEM6 significantly reduced tumour volume in a CRPC model of 22rv1-derived orthografts (assessed by ultrasonography, Fig. 1m). In this model, orchidectomy is performed at the time of the orthotopic cell transplantation to mimic the *in vivo* environment of ADT. To complement the use of 22rv1 orthografts to model CRPC, we further applied the CWR22res orthograft model to study the effects of acute ADT on pre-established tumours (orchidectomy 3 weeks post injection), thus better resembling treatment in clinical patients (24). Similar to 22rv1, THEM6 KO strongly impaired CRPC tumour growth in the CWR22res orthograft model (Fig. 1n). In addition to decreased tumour size, THEM6-deficient orthografts exhibited large necrotic areas and decreased cellularity, especially under ADT conditions (Suppl. Fig. 1j). Taken together, these data support a pro-tumorigenic role for THEM6 in PCa.

### THEM6 regulates cellular lipid metabolism

As a member of the THEM superfamily, THEM6 exhibits an evolutionarily conserved “Hotdog” domain predicted to confer thioesterase activity (18). Rewiring of lipid metabolism is a common feature of ADT resistance (15). Therefore, we hypothesised that THEM6 could participate in the lipid rearrangement required for CRPC progression. To evaluate the impact of THEM6 loss on the lipid composition of CRPC cells, we compared the lipid profiles of control (CTL) and THEM6 KO cells using LC-MS lipidomics. Strikingly, loss of THEM6 in 22rv1 cells resulted in a profound remodelling of the cellular lipidome. THEM6 depletion was associated with a strong reduction in the intracellular levels of multiple triglyceride (TG) and ether lipid species (ether triglycerides (ether TG) and ether phospholipids (ether PC or ether PE)). In contrast, THEM6 KO cells displayed increased amounts of ceramides (Fig. 2a). In addition to specific lipid changes, THEM6 KO also significantly affected the total amount of TGs, ether TGs and ceramides in 22rv1 cells (Fig. 2b). Interestingly, the intracellular levels of several ether lipid species (but not TG) were also strongly reduced in LNCaP AI THEM6 KO cells when compared to their respective CTL (Fig. 2c and Suppl. Fig. 2a), suggesting that THEM6 loss primarily affects ether lipid homeostasis. We further assessed the effect of THEM6 KO on lipid content *in vivo* by performing Raman spectroscopy on ADT-treated CWR22res orthografts. Raman spectroscopy allows assessment and quantitation of lipids (band at 2845 cm^-1^) and cholesterol (band at 2880 cm^-1^) content on paraffin-embedded tumour slides in a non-destructive manner (Fig. 2d). Results from the analysis provided evidence that THEM6-deficient tumours (Fig. 1n) display significantly less lipids (Fig. 2e) and cholesterol (Fig. 2f) than CTL tumours, thus confirming a role for THEM6 in the maintenance of the tumour lipidome.

**Figure 2:**
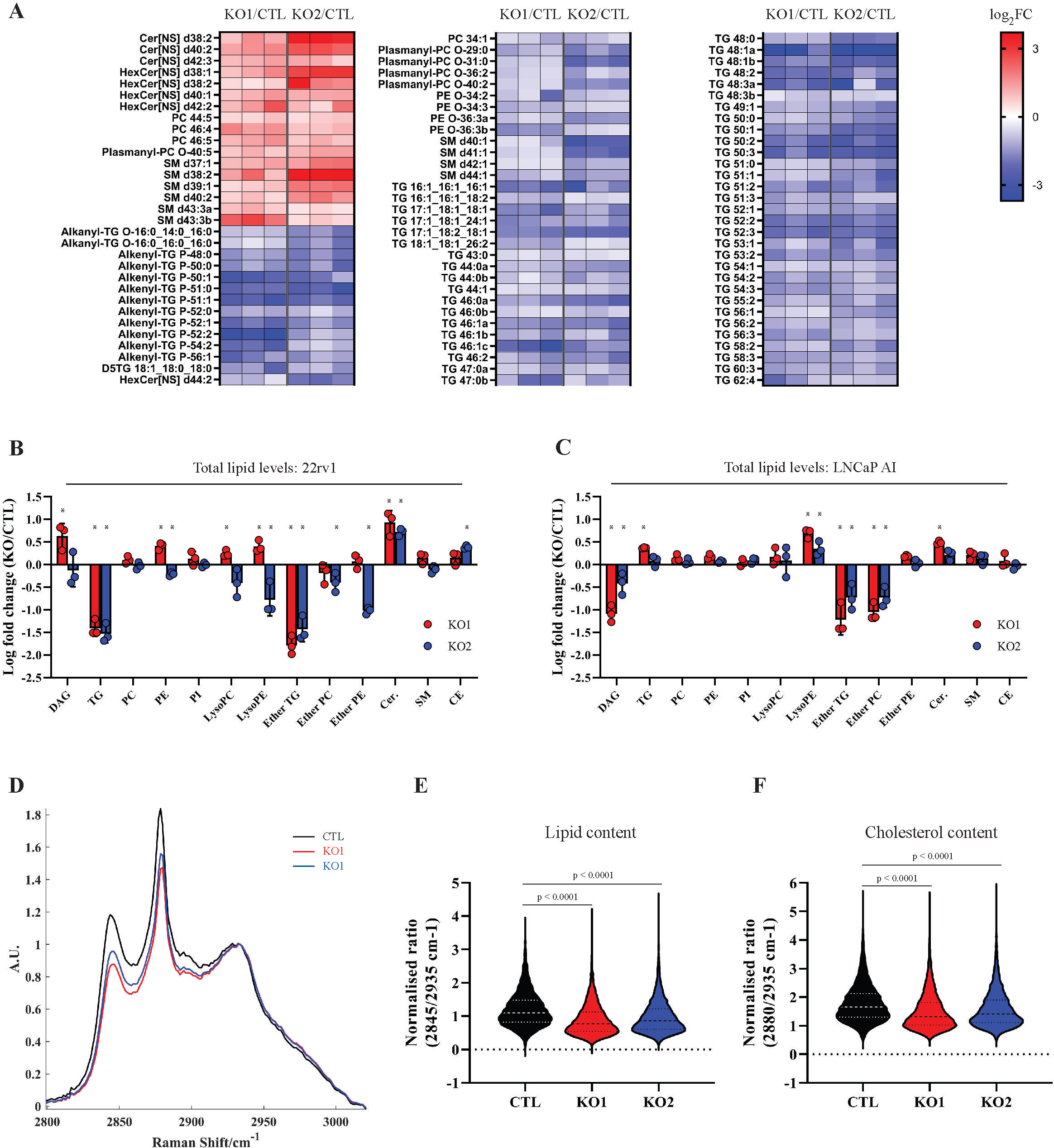
Loss of THEM6 alters lipid homeostasis. **A,** Heatmap illustrating the steady-state levels of significantly regulated lipids in THEM6 KO 22rv1 cells when compared to CTL (p ≤ 0.05, FC = 1.3). Values are expressed as log2(FC). **B-C,** Changes in lipid content (total amount) observed in THEM6 KO CRPC cells when compared to CTL. **D,** Average Raman spectra of CWR22res CTL and THEM6 KO orthografts. **E-F,** Quantification of tumour lipid (2845 cm^-1^-peak) and cholesterol (2880 cm^-1^-peak) content obtained from **D**. AU = arbitrary unit. Panels **B, C** : Data are presented as mean values +/- SD. Panels **E, F**: Centre line corresponds to median of data, top and bottom lines correspond to upper and lower quartiles. <u>Statistical analysis</u> : **B, C**: *p-value < 0.05 using a 1-way ANOVA with a Dunnett’s multiple comparisons test. **E, F**: Kruskal-Wallis test. DAG: diacylglycerol; TG: triglyceride; PC: phosphatidylcholine; PE: phosphatidylethanolamine; PI: phosphatidylinositol; LysoPC: lysophosphatidylcholine; LysoPE: lysophosphatidylethanolamine; Cer: Ceramide; SM: sphingomyelin; CE: Cholesteryl ester.

### THEM6 is an ER membrane protein that is essential to maintain ER integrity

Lipids are essential components of biological membranes. Interestingly, THEM6 depletion in CRPC cells resulted in increased cell stiffness (Fig. 3a), a phenotype which is indicative of membrane and cytoskeleton reorganisation and often associated with impaired migratory abilities (25). To gain further insights into the role of THEM6 in cancer cells, we performed a proteomic comparison of 22rv1 cells proficient or depleted for THEM6 (Fig. 3b). Proteomic analysis highlighted a large cluster of ER-related proteins that were significantly down-regulated in the absence of THEM6 (FC = 1.3, p < 0.05, Fig. 3c and Suppl. Table 1). In addition, the majority of these proteins are described as membrane proteins (Fig. 3c). ER is particularly sensitive to lipid perturbation (26). Electron microscopy confirmed the negative impact of THEM6 depletion on ER morphology. Indeed, THEM6 KO cells presented with abnormal ER, exhibiting rounded structures with highly dilated lumens (Fig. 3d). The presence of abnormally enlarged mitochondria and large multilamellar lysosomes was also frequently observed in THEM6 KO cells, potentially reflecting general membrane perturbations following THEM6 loss (Suppl. Fig. 3a). Interestingly, THEM6 strongly co-localised with the ER marker calreticulin (Fig. 3e, Suppl. Fig. 3b), but not with mitochondria (Suppl. Fig. 3c). This result suggests that THEM6 is predominantly associated with ER, contrasting with the mitochondrial localisation reported for other THEM superfamily members (THEM2/4/5) (27). Furthermore, topological analysis of the THEM6 protein sequence highlighted a 17-amino acids signal peptide that corresponds to a well-defined N-terminal transmembrane domain (Fig. 3f). Finally, western blot (WB) analysis after subcellular fractionation confirmed the presence of THEM6 in the insoluble organellular/membrane fraction of CRPC cells (Fig. 3g, Suppl. Fig. 3d).

**Figure 3:**
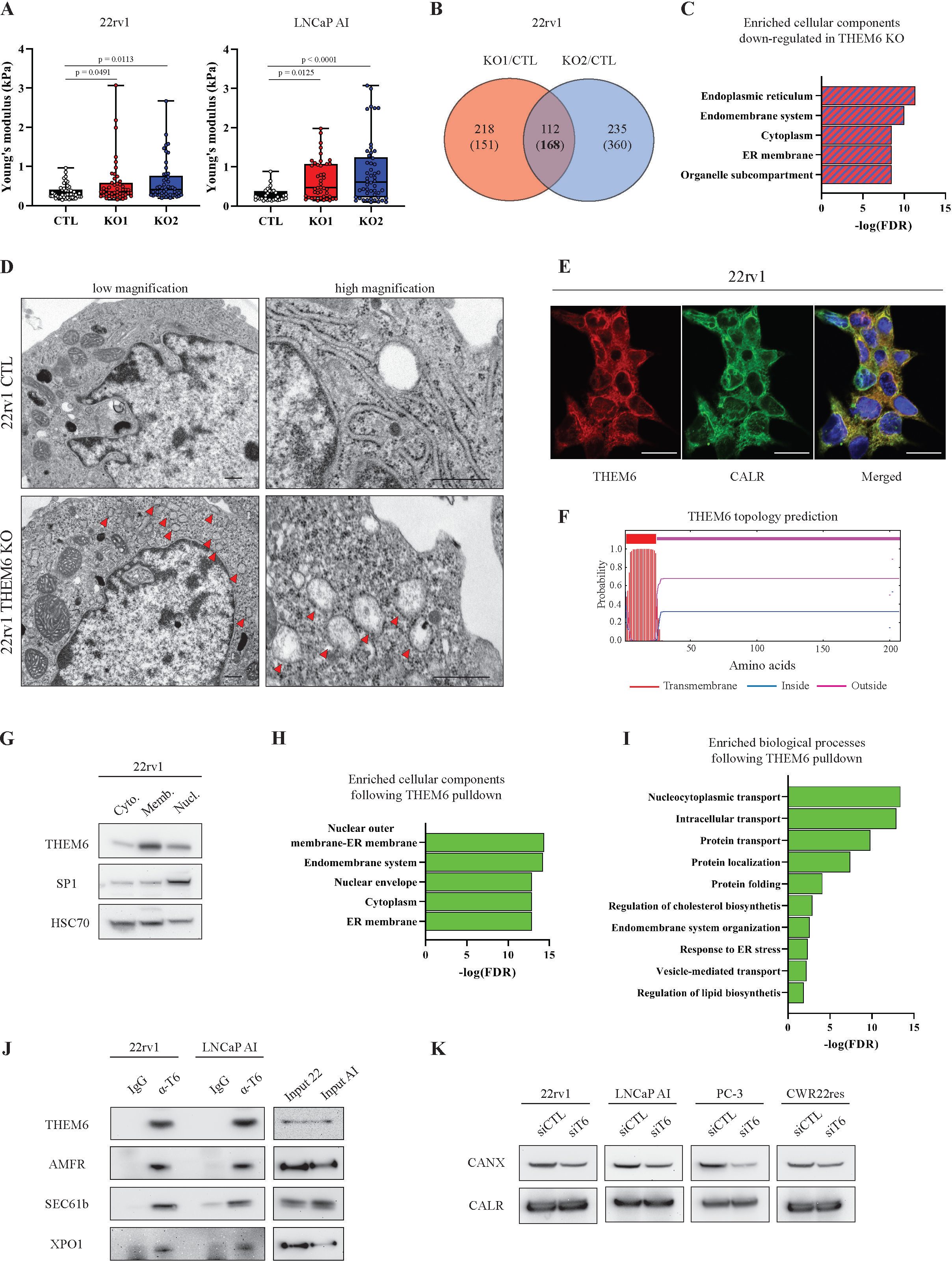
THEM6 regulates ER membrane trafficking. **A**, Cell stiffness (Young’s modulus) of CTL and THEM6 KO CRPC cells measured by atomic force microscopy. **B,** Venn diagram highlighting commonly modulated proteins (p-value ≤ 0.05, FC = 1.3) in THEM6 KO 22rv1 cells (2 clones) when compared to CTL. Up-regulated proteins are on top; Down-regulated proteins are into brackets. **C,** Enriched cellular components commonly down-regulated in THEM6 KO 22rv1 cells (2 clones) when compared to CTL. **D,** Representative electron microscopy pictures of CTL and THEM6 KO 22rv1 cells taken at low (left) and high (right) magnification. Red arrows point towards abnormal ER structure. Scale bar represents 500 nm. **E,** Immunofluorescence showing co-localisation of THEM6 and the ER marker calreticulin in 22rv1 cells. Scale bar represents 20 µm. **F,** Prediction of transmembrane domain in the sequence of the THEM6 protein. Topology prediction was performed using the TMHMM server (http://www.cbs.dtu.dk/services/TMHMM). **G**, Western blot analysis of THEM6 expression in cytoplasmic (cyto.), membrane/organelle (memb.) and nuclear fractions (nucl.) of 22rv1 cells. HSC70 and SP1 were used as cytoplasmic and nuclear-enriched markers, respectively. **H-I,** Enriched cellular components (**H**) and biological processes (**I**) in THEM6-interacting proteins following THEM6 pulldown in T6 OE HEK293 cells. **J**, Western blot analysis of THEM6, AMFR, SEC61b and XPO1 expression in CRPC cells following THEM6 immunoprecipitation in CRPC cells. **K**, Western blot analysis of CALX and CALR expression in PCa cells following THEM6 silencing. Panel **A**: Data are presented as mean values +/- SD. Panels **C, H, I**: Enrichment analysis were performed using the STRING database (http://string-db.org). <u>Statistical analysis</u> : **A**: 1-way ANOVA with a Dunnett’s multiple comparisons test.

### THEM6 regulates membrane protein trafficking in the ER

Lipid metabolism and protein homeostasis are highly interconnected processes (28). In particular, ether lipids and plasmalogen derivatives play a key role in regulating membrane protein trafficking and homeostasis (29). Due to its localisation at the ER membrane and its role in regulating lipid balance, THEM6 might therefore be important for the maintenance of protein homeostasis in the ER. In line with this idea, pull-down experiments followed by MS analysis identified 152 proteins that significantly interacted with THEM6 in THEM6-overexpressing HEK-293 cells (FC = 10, p < 0.05, Suppl. Fig. 3e and Suppl. Table 2). Most of the THEM6 interactors were located at the ER membrane, or at the interface between ER and nucleus (Fig. 3h), and were mainly involved in protein transport (Fig. 3i). Among others, several exportins, importins, transportins, as well as components of the oligosaccharyltransferase (OST) complex were identified as strong THEM6-interacting partners (Suppl. Table 2). Interactions between endogenous THEM6 and membrane proteins from different subcellular compartments (CRM1 at the outer nuclear membrane, AMFR and SEC61b in the ER) were confirmed using immunoprecipitation experiments in CRPC cell lines (Fig. 3j). Importantly, acute THEM6 silencing in PCa cell lines led to a consistent decrease in the expression of the ER membrane-associated lectin calnexin (CALX) but did not affect the levels of the soluble homolog calreticulin (CALR) (Fig. 3k), indicating that THEM6 loss might preferentially affect the trafficking of membrane proteins.

### THEM6 loss affects *de novo* sterol and FA synthesis in CRPC cells

In addition to protein trafficking, ER is also the primary site for lipid and cholesterol synthesis. Therefore, we postulated that THEM6 depletion would impact on these metabolic processes. Supporting this idea, our proteomic analysis of THEM6-deficient cells highlighted a down-regulation of multiple proteins involved in sterol biosynthesis (Fig. 4a). Sterol biosynthesis is of particular interest in the context of CRPC, as cholesterol serves as a precursor for *de novo* androgen synthesis and sustains ADT resistance (24, 30). We first validated the decreased expression of several enzymes involved in sterol biosynthesis in THEM6 KO cells (Fig. 4b). Next, we tested the effect of THEM6 loss on *de novo* sterol biosynthesis by incubating the cells with [U13C]-glucose and [U13C]-glutamine and following ^13^C incorporation into sterols using GC-MS. Surprisingly, we were not able to detect a significant proportion of labelled cholesterol in 22rv1 cells. Instead, these cells accumulated large amounts of *de novo* synthesised desmosterol, an immediate precursor of cholesterol (Fig. 4c and Suppl. Fig. 4a). In agreement with the proteomic data, 22rv1 THEM6 KO cells accumulated significantly less ^13^C-enriched desmosterol than CTL cells (Fig. 4c). In contrast to 22rv1 cells, the isogenic CWR22res cell line displayed a significant proportion of labelled cholesterol after incubation with [U13C]-glucose and [U13C]-glutamine. In this cell line, ^13^C-enrichment of cholesterol was also significantly reduced following THEM6 depletion (Fig. 4d and Suppl. Fig. 4b). Furthermore, using the PRAD TCGA dataset, we found that THEM6 expression strongly correlated with the expression of several enzymes involved in the late steps (SQLE, LSS, DHCR7 and DHCR24, Fig. 4e), but not in the early steps (mevalonate pathway, Suppl. Fig. 4c), of sterol biosynthesis, suggesting that THEM6 also contributes to this pathway in PCa patients. Similarly, unbiased analysis of the PRAD TCGA dataset revealed that Acetyl-CoA Carboxylase (ACACA) and Fatty Acid Synthase (FASN), two regulatory enzymes of the fatty acid (FA) synthesis, were among the top up-regulated proteins in THEM6-enriched patient tumours (Fig. 4f and Suppl. Table 3). We then assessed the contribution of THEM6 to FA synthesis by determining the relative proportions of ^13^C-labelled palmitic, oleic and stearic acids in presence and absence of THEM6 (Fig. 4g and Suppl. Fig. 4d). Similar to sterols, THEM6 KO cells accumulated significantly less labelled FA when compared to CTL, indicating reduced FA synthesis in absence of THEM6. Altogether, these results suggest that THEM6 expression is critical for the regulation of *de novo* lipid synthesis.

**Figure 4:**
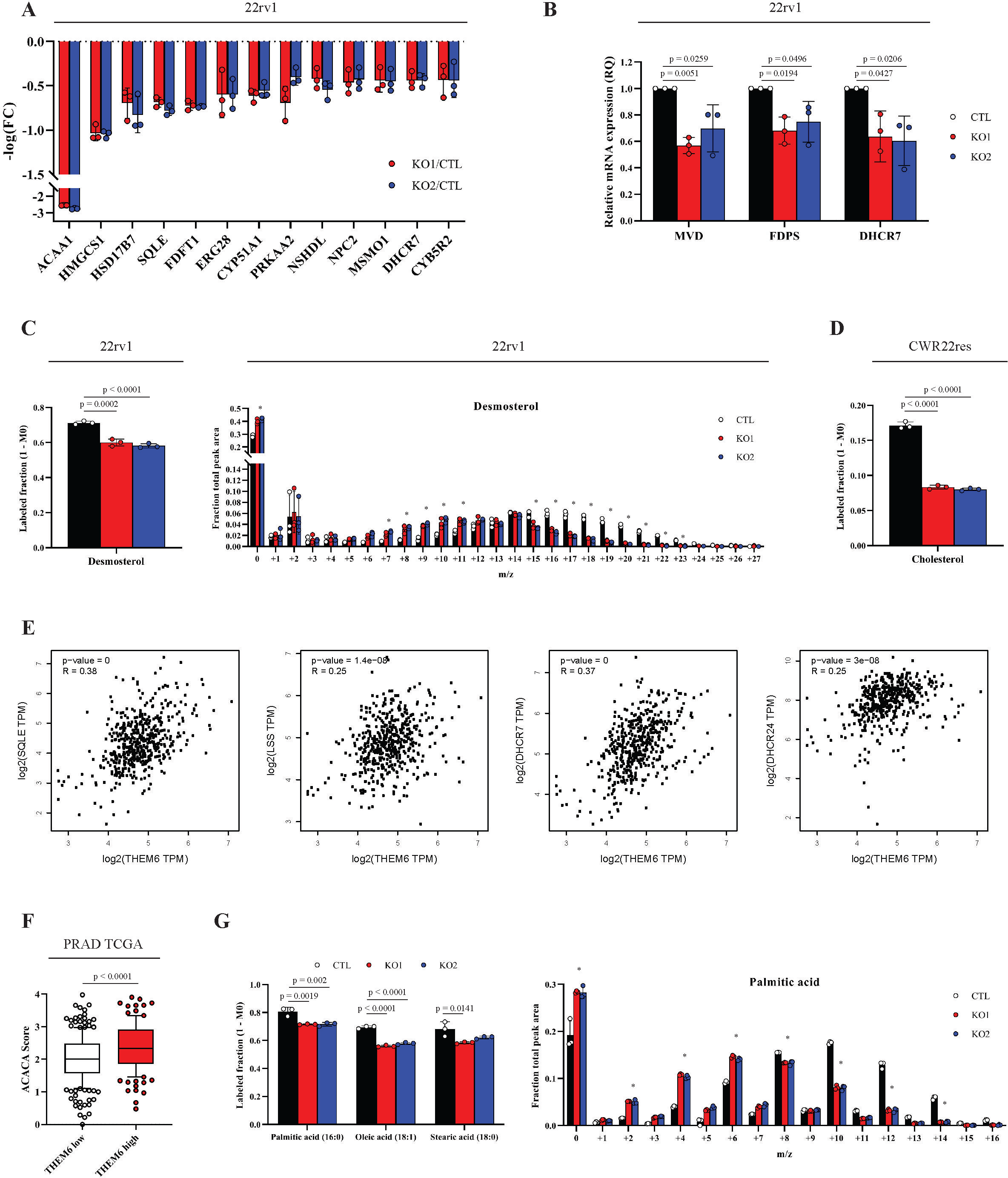
Loss of THEM6 affects de novo sterol and fatty acid synthesis. **A,** Proteomic analysis highlighting proteins associated with sterol biosynthetic pathway and significantly down-regulated in THEM6 KO 22rv1 cells when compared to CTL (p-value ≤ 0.05, FC = 1.3). **B,** RT-qPCR analysis of *MVD, FDPS* and *DHCR7* expression in CTL and THEM6 KO 22rv1 cells. *CASC3* was used as a normalising control. **C,** Labelled desmosterol fraction derived from ^13^C-glucose and ^13^C-glutamine (left panel) and relative isotopologue distribution of desmosterol in CTL and THEM6 KO 22rv1 cells after 72 hours of incubation (right panel). **D,** Labelled cholesterol fraction derived from ^13^C-glucose and ^13^C-glutamine in CTL and THEM6 KO CWR22res cells after 72 hours of incubation. **E,** Pearson’s correlation analysis of *SQLE*, *LSS, DHCR7* and *DHCR24* with *THEM6* using the PRAD TCGA dataset. Results were obtained using the GEPIA website http://gepia.cancer-pku.cn/. **F**, Differential expression of ACACA in high and low THEM6 tumours according to the PRAD TCGA dataset. **G,** Labelled palmitic, oleic and stearic acid fractions derived from ^13^C-glucose and ^13^C-glutamine (left panel) and relative isotopologue distribution of palmitic acid in CTL and THEM6 KO 22rv1 cells after 72 hours of incubation (right panel). Panels **A, B, C, D, G**: Data are presented as mean values +/- SD. Panel **F**: Centre line corresponds to median of data, top and bottom of box correspond to 90th and 10th percentile, respectively. Whiskers extend to adjacent values (minimum and maximum data points not considered outliers). <u>Statistical analysis</u> : **B, C, D, G**: *p-value < 0.05 using 1-way ANOVA with a Dunnett’s multiple comparisons test. **F**: two-tailed Mann-Whitney U test.

### THEM6 is required to trigger ATF4 induction in CRPC cells

Perturbation of ER homeostasis results in the activation of a tightly regulated stress response program, the unfolded protein response (UPR), in order to rapidly alleviate ER stress (31). Enrichment pathway analysis of the THEM6-deficient 22rv1 cells highlighted “Response to ER stress” as the main pathway regulated in absence of THEM6 (Fig. 5a), with 17 proteins referenced in this pathway significantly down-regulated in both KO clones (FC = 1.3, p < 0.05, Fig. 5b and Suppl. Table 1). Interestingly, BIP (HSPA5), the main regulator of the UPR, was identified among the proteins significantly down-regulated in 22rv1 THEM6 KO cells (Fig. 5b), and we confirmed this result by WB (Fig. 5c). Impaired UPR activation in THEM6-deficient cells was further evidenced by decreased levels of the UPR effectors XBP1s (spliced isoform of XBP1), ATF4 and CHOP (DDIT3) (Fig. 5c). As a consequence, THEM6 KO significantly sensitised CRPC cells to prolonged ER stress, caused by chronic tunicamycin treatment (Fig. 5d).

**Figure 5:**
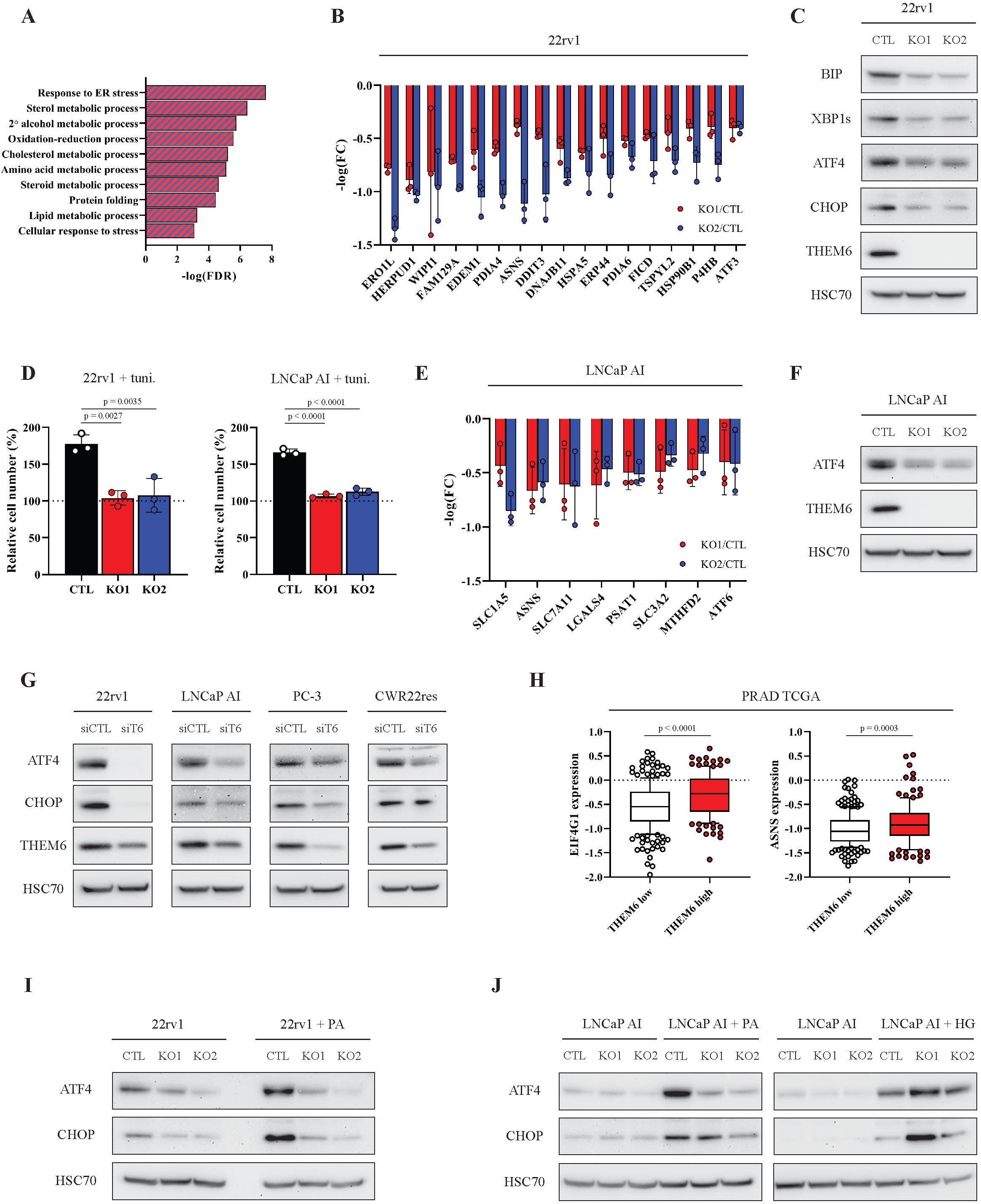
THEM6-mediated lipid remodelling is required for UPR activation. **A,** Enriched biological processes commonly down-regulated in THEM6 KO 22rv1 cells (2 clones) when compared to CTL. **B,** Proteomic analysis highlighting proteins associated with the ER stress response significantly down-regulated in THEM6 KO 22rv1 cells when compared to CTL (p-value ≤ 0.05, FC = 1.3). **C**, Western blot analysis of BIP, XBP1s, ATF4, CHOP and THEM6 expression in CTL and THEM6 KO 22rv1 cells. **D**, Proliferation of CTL and THEM6 KO CRPC cells treated with tunicamycin (2.5 µg/ml) for 72 hours. Cell count is normalised to initial number of cells at T0. **E,** Proteomic analysis highlighting ATF4 targets significantly down-regulated in THEM6 KO LNCaP AI cells when compared to CTL (p-value ≤ 0.05, FC = 1.3). **F**, Western blot analysis of ATF4 and THEM6 expression in CTL and THEM6 KO LNCaP AI cells. **G**, Western blot analysis of ATF4, CHOP and THEM6 expression in PCa cells following THEM6 silencing. **H**, Differential expression of EIF4G1 and ASNS in high and low THEM6 tumours according to the PRAD TCGA dataset. **I**, Western blot analysis of ATF4 and CHOP expression in CTL and THEM6 KO 22rv1 cells treated with palmitic acid (200 µM) for 48 hours. **J**, Western blot analysis of ATF4 and CHOP expression in CTL and THEM6 KO LNCaP AI cells treated with palmitic acid (200 µM) or hexadecylglycerol (50 µM) for 48 hours. Panels **C, F, G, I, J**: HSC70 was used as a sample loading control. Panels **B, D, E**: Data are presented as mean values +/- SD. Panel **H**: Centre line corresponds to median of data, top and bottom of box correspond to 90th and 10th percentile, respectively. Whiskers extend to adjacent values (minimum and maximum data points not considered outliers). <u>Statistical analysis</u> : **D**: 1-way ANOVA with a Dunnett’s multiple comparisons test. **H**: two-tailed Mann-Whitney U test.

Proteomic analysis of THEM6 KO in LNCaP AI cells also highlighted ER perturbation as the main consequence of THEM6 depletion (Suppl. Fig. 5a and Suppl. Table 4). Of note, THEM6 KO in this cell type led to an accumulation of several ER chaperones, including BIP, and did not consistently affect XBP1s levels (Suppl. Fig. 5b). In contrast with IRE1a-XBP1s signalling, proteomic analysis highlighted many ATF4 targets that were down-regulated in LNCaP AI THEM6 KO cells (Fig. 5e), suggesting that THEM6 might be particularly important for ATF4 activation in PCa. In line with this idea, we found that ATF4 levels were strongly reduced in THEM6-deficient LNCaP AI cells (Fig. 5f), and that both ATF4 and CHOP were also decreased following transient THEM6 silencing in multiple PCa cell lines (Fig. 5g). In addition, analysis of the PRAD TCGA data revealed that EIF4G1, a member of the EIF4F complex required for ER stress-induced translation of ATF4, and ASNS, a canonical ATF4 target, were significantly enriched in tumours with high THEM6 expression (Fig. 5h and Suppl. Table 3), thus underpinning the importance of THEM6 in the establishment of the ATF4-mediated stress response.

Lipid-mediated stress leads to UPR activation (26). Because of the implication of THEM6 in maintaining lipid homeostasis (Fig. 2 and Fig. 4), we wondered whether the inability of the THEM6-deficient cells to induce ATF4 was due to changes in their lipid composition. While CTL cells strongly induced ATF4 expression following palmitic acid (PA) treatment, ATF4 levels remained unaffected by PA in THEM6 KO cells (Fig. 5i). Importantly, treatment with the ether lipid precursor hexadecylglycerol (HG) induced ATF4 expression to a similar level in both THEM6 KO and CTL cells (Fig. 5j). Taken together, these results suggest that the stress-induced activation of ATF4 depends, at least in part, on THEM6-mediated lipid remodelling in CRPC.

### THEM6 is required for MYC-dependent activation of ATF4

ADT is a major source of metabolic stress (32). In PCa, MYC is an important regulator of the UPR and promotes tumour survival in androgen-independent conditions (9, 33). Importantly, at gene level, we observed that *THEM6* was located alongside *MYC* on the chromosome locus 8q24.3, a genomic region that is frequently dysregulated in cancer (34). Consequently, we found that *THEM6* was co-amplified with *MYC* in multiple prostate cancer datasets (Fig. 6a-b). On average *THEM6* and *MYC* genomic alterations occurred in 34% and 38% of PCa patients, and this incidence was even higher in metastatic cases (average genomic alteration frequency of 51% and 56% for *THEM6* and *MYC* respectively, Fig. 6a). Accordingly, MYC expression was high in CRPC tumour orthografts compared to their HN counterparts (Fig. 6c), and thus correlated with THEM6 expression (Fig. 1b). Furthermore, acute ADT was sufficient to increase both THEM6 and MYC levels in the CWR22Res orthograft model (Suppl. Fig. 6a). To evaluate the importance of THEM6 function in the context of high MYC signalling, we used mouse embryonic fibroblasts (MEFs) as well as U2OS cells expressing a tamoxifen-inducible MYC-ER construct, hereafter referred to as MEFs-MYC^ER^ and U2OS-MYC^ER^ respectively. MYC activation in MEFs-MYC^ER^ strongly increased THEM6 expression (Fig. 6d), and transient silencing of THEM6 significantly impaired the MYC-induced proliferation of these cells (Fig. 6e). Of note, prolonged MYC activation resulted in a strong accumulation of the THEM6 protein over time (Suppl. Fig. 6b) while its increased expression was transient at the mRNA level (∼3-fold after 4h to 6h, Suppl. Fig. 6c), indicating that additional regulations might occur at the protein level. Activation of MYC in U2OS-MYC^ER^ increased THEM6 protein expression and further led to a strong accumulation of ATF4 and CHOP (Fig. 6f). Strikingly, transient THEM6 silencing was sufficient to abolish the MYC-induced expression of these two factors, as well as strongly reducing CALX levels (Fig. 6f). In addition, THEM6 depletion significantly altered the lipid profile of MYC-activated cells, resulting in decreased levels of many ether PCs and PEs (Fig. 6g). Altogether, these results suggest that THEM6 is essential for the establishment of a MYC-dependent stress response.

**Figure 6:**
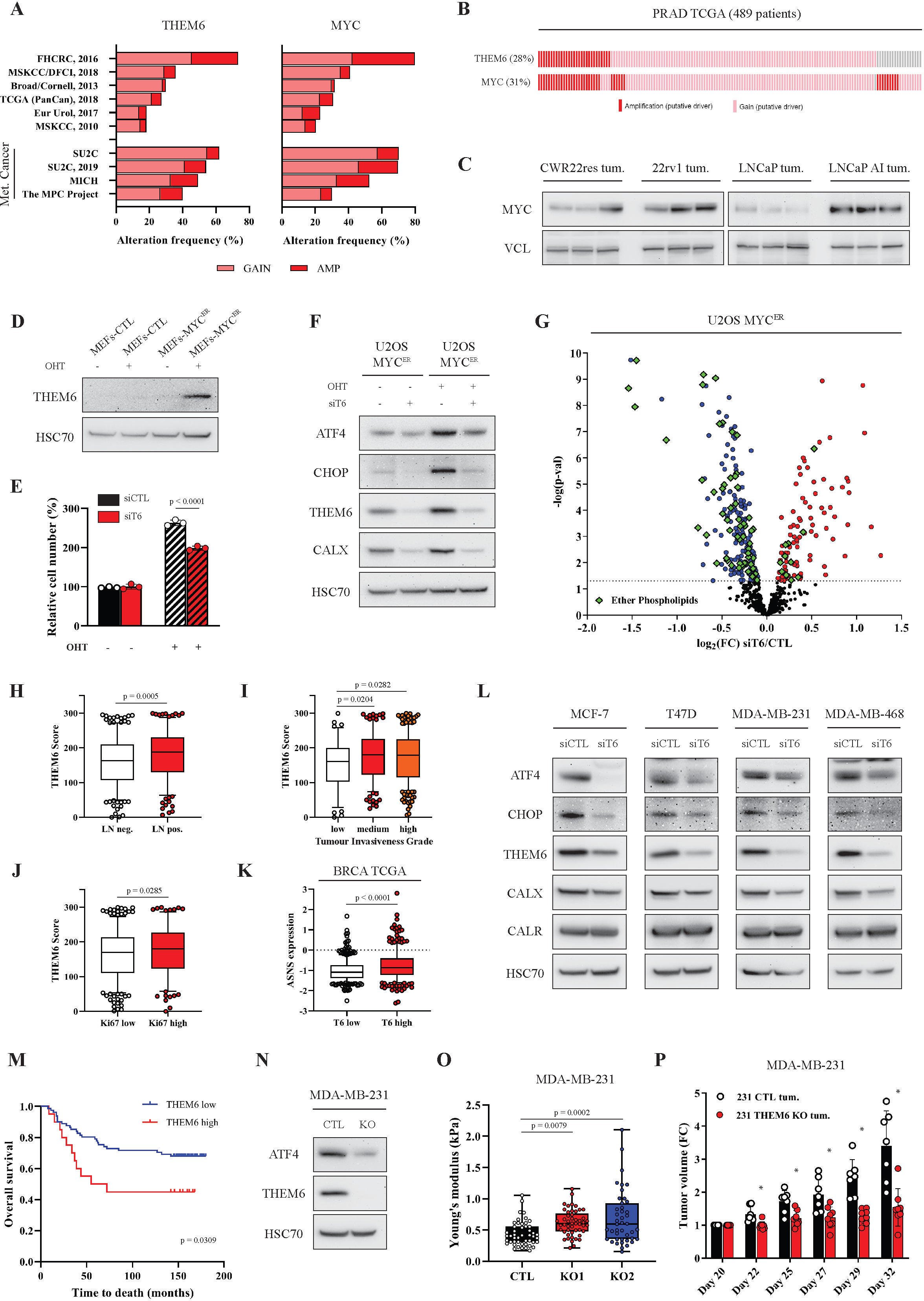
THEM6 is an actionable target in high MYC cancer. **A**, Percentage of PCa patients showing genomic alteration (copy number gain or amplification) for *THEM6* (left) and *MYC* (right) in publicly available datasets of PCa. Analysis was performed using the cbioportal website https://www.cbioportal.org/. **B**, Oncoprint analysis showing the percentage of PCa patients with co-occuring *THEM6* and *MYC* genomic alterations (copy number gain or amplification) in the PRAD TCGA. Analysis was performed using the cbioportal website https://www.cbioportal.org/. **C**, Western blot analysis of MYC expression in CRPC (22rv1 and LNCaP AI) and HN (CWR22res and LNCaP) prostate orthografts. **D**, Western blot analysis of THEM6 expression in CTL and MYC-inducible MEFs. **E**, Proliferation of CTL and THEM6 KD MEFs following MYC induction for 48 hours. **F**, Western blot analysis of ATF4, CHOP, CALX and THEM6 expression in CTL and MYC-inducible U2OS cells. **G,** Volcano plot of the differentially modulated lipids in THEM6 KD OHT-treated U2OS cells when compared to OHT-treated CTL (p ≤ 0.05). Red and blue dots represent the lipids that are significantly up- and down-regulated, respectively. Ether phospholipids are highlighted in green. **H-J**, Quantification of THEM6 expression in BCa tissue samples according to lymph node positivity (**H**), tumour invasiveness (**I**) and Ki67 score (**J**). **K**, Differential expression of ASNS in high and low THEM6 tumours according to the BRCA TCGA dataset. **L**, Western blot analysis of ATF4, CHOP, THEM6, CALX and CALR expression in BCa cells following THEM6 silencing. **M**, Kaplan-Meier overall survival analysis of TNBC patients stratified according to upper quartile THEM6 expression. **N**, Western blot analysis of ATF4 expression in CTL and THEM6 KO MDA-MB-231 cells. **O**, Cell stiffness (Young’s modulus) of CTL and THEM6 KO MDA-MB-231 cells measured by atomic force microscopy. **P**, Tumour volume of CTL and THEM6 KO MDA-MB-231-derived orthografts. Panels **C, D, F, L, N**: HSC70 was used as a sample loading control. Panels **E, O, P**: Data are presented as mean values +/- SD. Panels **H, I, J, K**: Centre line corresponds to median of data, top and bottom of box correspond to 90th and 10th percentile, respectively. Whiskers extend to adjacent values (minimum and maximum data points not considered outliers). <u>Statistical analysis</u> : **E**: two-tailed Student’s t-test. **H, J, K, P**: two-tailed Mann-Whitney U test. **I**: Kruskal-Wallis test. **M**: logrank test. **O**: 1-way ANOVA with a Dunnett’s multiple comparisons test.

### THEM6 is an actionable target in multiple tumour types

In addition to genomic amplification, THEM6 overexpression has been reported in multiple cancer types (35). Moreover, the ability of THEM6 to facilitate MYC-induced UPR suggests that the pro-tumoural effects of THEM6 might not be restricted to prostate cancer. Meta-analysis of THEM6 expression across the different TCGA datasets revealed that THEM6 overexpression occurred frequently in hormone-dependent (Suppl. Fig. 6d) as well as liver-and gastrointestinal cancers (Suppl. Fig. 6e). We further investigated THEM6 in breast tumours where hormone therapy is also routinely used as primary cancer treatment strategy. Using a breast cancer TMA (n = 551), we found that high THEM6 levels were associated with poorer clinical parameters, such as enhanced lymph node infiltration (Fig. 6h), increased tumour invasiveness (Fig. 6i) and high levels of Ki67 (Fig. 6j). Similar to what we observed in PCa (Fig. 5h), high THEM6 expression correlated with high ASNS expression in the BRCA TCGA dataset (Fig. 6k), and THEM6 silencing led to reduced levels of ATF4, CHOP and CALX in multiple breast cancer cell lines (Fig. 6l).

Breast cancer patients with high THEM6 expression further displayed a trend towards reduced survival (Suppl. Fig. 6f). Strikingly, this effect was emphasised in triple negative breast cancer (TNBC) (n = 123, Fig. 6m), an aggressive molecular subtype that heavily relies on MYC expression (36). Therefore, in a proof-of-concept experiment, we assessed the impact of THEM6 loss in TNBC cells. Consistent with a role for THEM6 in UPR regulation, THEM6 KO in MDA-MB-231 cells led to a strong decrease in ATF4 expression (Fig. 6n). Finally, similarly to the PCa, loss of THEM6 increased cell stiffness (Fig. 6o) and impaired *in vivo* tumour growth in the TNBC model (Fig. 6p, Suppl. Fig. 6g), thus supporting a general pro-tumoural role for THEM6 in cancer.

## DISCUSSION

The remarkable reliance of PCa on AR signalling led to the generation of effective targeted therapies, such as enzalutamide, abiraterone acetate and other small molecule inhibitors of the AR pathway, which have significantly improved the clinical management of non-resectable prostate tumours. However, resistance to such therapies ultimately results in the development of lethal CRPC. A detailed characterisation of the molecular signalling pathways associated with treatment resistance will therefore identify novel actionable targets and foster innovative therapeutic options. For example, a better understanding of treatment-induced metabolic rewiring (15) or specific dependencies on stress response pathways (5) might uncover vulnerabilities with the potential to synergise with current treatments. Our lab recently engaged in a comparison of different *in vivo* models of ADT resistance at multi-omics levels (21). From this comprehensive dataset, we identified Thioesterase Superfamily Member 6 (THEM6) as a clinically relevant protein overexpressed in CRPC. In patients, THEM6 expression correlated with tumour aggressiveness and was highest in treatment-resistant tumours. Furthermore, high THEM6 expression was associated with poor clinical tumour parameters, such as enhanced tumour proliferation and metastatic dissemination, and correlated with shortened overall and disease-free patient survival. Importantly, the prognostic potential of THEM6 was not restricted to PCa but could also be applicable to other cancer types, such as TNBC, which at present have few treatment options. THEM6, formerly c8orf55, was originally discovered as a potential biomarker for colon cancer in a proteomic-based analysis of human samples. The authors further validated THEM6 overexpression in several tumour types, including breast and prostate (35). THEM6 has also been reported in additional proteomic studies, mainly comparing tumoural and respective non-malignant tissues from various origins (37, 38).

Despite no characterised biological function, THEM6 has recently been classified in the THEM thioesterase superfamily due to the presence of a “HotDog” domain, an evolutionarily conserved domain with a predicted thioesterase activity (17). Members of the THEM superfamily (THEM1/ACOT11, THEM2/ACOT13, THEM4 and THEM5) display an acyl-CoA thioesterase activity, although presenting differences in specificity towards their fatty acyl-CoA substrates. Consequently, the THEM proteins have been described to play various roles in FA and lipid metabolism (18). In line with these studies, we found that THEM6 depletion significantly altered the lipid composition of cancer cells. In particular, loss of THEM6 resulted in consistent decreases in the levels of several ether lipid classes (ether TGs and ether PCs). Despite being relatively understudied in this context, ether lipids are frequently over-represented in tumours (39) and support a pro-oncogenic phenotype (40). Ether lipids are a subset of glycerolipids characterised by the presence of an ether-alkyl or a vinyl ether-alkyl (plasmalogens) bond at the glycerol sn-1 position. Their synthesis is initiated in peroxisomes and terminates in the ER, where they undergo active acyl chain remodelling (41). In addition to their roles in redox homeostasis (42, 43) and cellular signalling, ether lipids are essential components of biological membranes. The importance of ether lipids in membrane protein trafficking, as well as their coordinate regulation with sphingolipids, has recently been described in mammalian cells (44). Interestingly, we identified THEM6 as an ER membrane protein whose absence primarily impacts ER function and morphology. Indeed, THEM6-deficient cells displayed abnormal ER structure with the presence of highly dilated lumen, a phenotype that is also observed following dysregulation of ether lipid metabolism (45). The observation that both mitochondrial and lysosomal structures were also affected in THEM6 KO cells further suggests that THEM6 loss induces a global perturbation of membrane homeostasis. Moreover, THEM6 closely interacted with many membrane proteins involved in protein transport (exportins, importins, transportins, components of the ERAD machinery and the OST complex), and THEM6 silencing led to a selective down-regulation of the membrane-associated chaperone calnexin, without affecting its soluble homolog calreticulin. CALX and CALR share substantial sequence identity, but the former is a type 1 transmembrane protein while the second mainly resides in the ER lumen (46). Of note, this decrease in CALX levels was much more pronounced upon transient siRNA transfection than following CRISPR-mediated KO, suggesting that THEM6-deficient cells might have adapted to compensate for this effect. Taken together, these results suggest that THEM6 is involved in the regulation of membrane homeostasis and trafficking. An appealing hypothesis would be that THEM6 influences membrane properties by controlling the balance between ester- and ether-bound lipids within the ER, for example by regulating the pool of FA that must be incorporated into ether lipids. The fact that THEM6 is the only member of the THEM superfamily presenting with a transmembrane domain supports this hypothesis, but at the same time renders the purification and structural analysis of the THEM6 protein very challenging. Therefore, additional work remains to be done to fully characterise THEM6 enzymatic activity and uncover its biochemical substrates.

ER is essential for the maintenance of both lipid and protein homeostasis. Perturbation of ER homeostasis, induced by increased proteo- or lipotoxicity, activates a complex signalling network, called the UPR, which aims at rapidly reducing ER stress (31). Importantly, we demonstrate that THEM6 is required for a correct activation of the UPR. In particular, the inability of cancer cells to activate the ATF4/CHOP pathway was a constant feature of THEM6-deficient cancer cells, irrespective of their origin. Furthermore, THEM6-depleted cells were unable to respond to palmitate-induced ER stress, but managed to activate the UPR in response to treatment with hexadecylglycerol, a precursor of the ether lipid synthesis. This result not only demonstrates the implication of THEM6 in ether lipid metabolism, but also highlights its importance in the establishment of the lipid-mediated stress response.

Importantly, failure to activate the UPR in response to ER stress has also been reported in the context of THEM1 and THEM2 deficiencies. Indeed, genetic ablation of *Them1* in mice attenuated diet- and chemical-induced ER stress responses, as well as UPR-mediated lipogenesis (19). Of note, the authors also speculated that *Them1* could influence ER stress levels by modulating the phospholipid composition of the ER membrane. Similarly, an elegant study from the same group demonstrated that *Them2*-mediated trafficking of saturated fatty acid critically regulates ER membrane fluidity, and is therefore required for calcium-dependent induction of ER stress in non-physiological condition (20).

ER stress induction and subsequent activation of the UPR have been described to promote tumorigenesis and treatment resistance in cancer (47). In PCa, resistance to ADT involves the activation of a sustained stress response together with major metabolic reprogramming (15, 48). An important contributor to these two processes is the MYC oncogene, which promotes androgen-independent growth and survival in PCa cells (9). *MYC* amplification is frequent in advanced PCa and correlates with enhanced tumour aggressiveness and increased resistance to treatment (49). Concomitant activation of MYC and the UPR is commonly observed in cancer, and both processes are tightly interconnected (33). As such, therapeutic targeting of the UPR has emerged as a promising strategy for the treatment of MYC-reliant tumours, including prostate (2,6–8,10). Interestingly, *THEM6* is co-amplified with *MYC* in cancer patient biopsies, while THEM6 expression is consistently increased upon MYC activation in different cell models. Furthermore, we showed that THEM6 is required for the correct activation of ATF4 following MYC induction. In addition to its role in cancer metabolism (50), ATF4 is essential for the regulation of MYC-dependent translation in cancer (51). Moreover, ATF4 has also been highlighted as an important regulator of PCa growth and survival (11). Along the same lines, Nguyen et al. demonstrated that the PERK-eiF2α pathway is selectively activated in metastatic PCa, and that targeting of this pathway significantly impairs tumour growth and metastasis dissemination in preclinical models (10). Overall, targeting oncogene-induced cellular or metabolic dependencies represents a promising approach for the treatment of aggressive cancers (6, 52). Therefore, through its role in lipid remodelling and its ability to regulate oncogene-induced UPR, THEM6 might provide an actionable target with the potential to be considered for the development of future anticancer therapies.

## METHODS

### Cell culture

LNCaP, C4-2, CWR22res, VCaP, DU-145, PC-3 and PC-3met were cultured in RPMI (Gibco, Thermo Fisher Scientific, Waltham, MA, USA) medium supplemented with 10% foetal bovine serum (Gibco, Thermo Fisher Scientific, Waltham, MA, USA) and 2 mM glutamine (Gibco, Thermo Fisher Scientific, Waltham, MA, USA). 22rv1 and LNCaP AI cells were maintained in phenol-free RPMI (Gibco, Thermo Fisher Scientific, Waltham, MA, USA) supplemented with 10% charcoal-stripped serum (Gibco, Thermo Fisher Scientific, Waltham, MA, USA) and 2 mM glutamine. MDA-MB-231, MCF-7, T47D, MDA-MB-468, MEFs-MYC^ER^ and U2OS-MYC^ER^ were maintained in DMEM (Gibco, Thermo Fisher Scientific, Waltham, MA, USA) supplemented with 10% foetal bovine serum (Gibco, Thermo Fisher Scientific, Waltham, MA, USA) and 2 mM glutamine. CWR22Res cells (hormone-responsive variant of CWR22 cells) were obtained from Case Western Reserve University, Cleveland, Ohio. MEFs expressing MYC^ER^ were isolated from Rosa26-MycER^T2^ embryos (53). U2OS-MYC^ER^ cells were generated in the lab of Daniel Murphy by transfection of U2OS cells with pBabe-MycER^T2^, followed by selection on puromycin. All other cell lines were obtained from ATCC. All cells were maintained at 37°C under 5% CO2 and routinely harvested with trypsin.

### Generation of CRISPR THEM6 KO cells

1×10^6^ cells were transfected with commercially available THEM6/DSCD75 KO or CTL plasmids (Santa Cruz Technologies, Dallas, TX, USA) using a nucleofector (Amaxa Biosystems, Lonza, Basel, Switzerland) according to the manufacturer’s instructions. Clonal selection of THEM6 KO cells was performed using puromycin (Sigma Aldrich, St Louis, MI, USA). Protein depletion was confirmed using western blot.

### Cell proliferation

1×10^6^ cells were seeded in 6-well plates and allowed to attach overnight. The next day, cells were harvested with trypsin (T0) or allowed to grow for additional 72 hours. Cells were then counted using a CASY cell counter (Roche, Basel, Switzerland). Final cell number was normalised to the initial cell count obtained at T0. Data are expressed as relative percentage of CTL cells.

### Measurement of cell stiffness using Atomic Force Microscopy

The mechanical properties of individual cells were measured using an Atomic Force Microscope Nanowizard II (JPK Instruments, Bruker, Berlin, Germany) with cell heater attachment mounted on an inverted optical microscope (Zeiss Observer Axio A.1, Zeiss, Cambridge, UK). Force indentation measurements were carried out as described previously (25, 54). Briefly, the AFM colloidal probes were prepared by gluing a 5.2 µm spherical silica bead (Bangs Laboratories, Inc, Fishers, IN, USA) at the end of a tipless cantilever (Arrow-TL2-50 with a spring constant of 0.03 N m^-1^, Nanoworld Innovative Technologies, Neuchatel, Switzerland) and calibrated prior to use. Cells were seeded on petri dishes at a concentration of 10^4^ cells cm^-2^ and cultured overnight. During the experiment, cells were kept at 37°C and in 25 mM HEPES buffered medium to maintain pH levels. Five points over the central nucleus area of each cell were measured, with ∼50 cells per population taken from multiple dishes to ensure reproducibility. The Young’s modulus was derived by fitting the extension part of a force–distance curve with the Hertzian spherical model using an in-house R Script (55). GraphPad Prism 8.4.2 was used to produce graphs and perform statistical analysis of the data.

### Proteomic analysis

1×10^6^ cells were seeded in 6-well plates and allowed to grow for 48 h. Cells were then lysed in SDS-containing buffer (2% SDS, 50 mM triethylammonium bicarbonate, pH 8.5) and briefly sonicated and centrifuged at 16,000 × g for 5 min at 4 °C. Protein concentration was determined using BCA assay (Thermo Fisher Scientific, Waltham, MA, USA) and samples were stored at −80 °C until further processing. For each experiment, 40 µg of protein extracts were reduced with 10 mM DTT and newly generated thiols were subsequently alkylated with 55 mM iodoacetamide. Alkylated proteins were precipitated using trichloroacetic acid (TCA). Washed pellets were reconstituted in 50 µl of 200 mM HEPES and digested first with Endoproteinase Lys-C (ratio 1:33 enzyme:lysate, Alpha Laboratories, Eastleigh, UK) for one hour, followed by an overnight trypsin digestion (ratio 1:33 enzyme:lysate, Promega, Madison, WI, USA).

The digested peptides from each experiment, and a pool sample, were differentially labelled using TMT10-plex reagent (Thermo Fisher Scientific, Waltham, MA, USA). Each sample was labelled with 0.1 mg of TMT reagent dissolved in 50 μl of 100% anhydrous acetonitrile. The reaction was carried out at room temperature for 2 hours. Fully labelled samples were mixed in equal amount and desalted using a Sep Pak C18 reverse phase solid-phase extraction cartridges (Waters corporation, Milford, MA, USA). After clean-up, the TMT-labelled peptides were fractionated using high pH reverse phase chromatography on a C18 column (150 × 2.1 mm i.d. - Kinetex EVO (5 μm, 100 Å)) using an HPLC system (Agilent, LC 1260 Infinity II, Agilent, Santa Clara, CA, USA). A two-step gradient was applied, from 1– 28% B in 42 min, then from 28–46% B in 13 min to obtain a total of 21 fractions for MS analysis.

Pull-down experiments were performed on Myc-tagged THEM6 overexpressing cells (hereafter referred to as Affinity Purification Mass Spectrometry experiment, AP-MS). Agarose beads were resuspended in a 2 M Urea and 100 mM ammonium bicarbonate buffer and stored at -20°C. Biological triplicates were digested with Lys-C (Alpha Laboratories, Eastleigh, UK) and trypsin (Promega, Madison, WI, USA) on beads as previously described (56). Prior mass spectrometry analysis, digested peptides were desalted using StageTip (57).

Peptides from all experiments were separated by nanoscale C18 reverse-phase liquid chromatography using an EASY-nLC II 1200 (Thermo Fisher Scientific, Waltham, MA, USA), using a binary gradient with buffer A (2% acetonitrile) and B (80% acetonitrile), both containing 0.1% formic acid. Samples were loaded into fused silica emitter (New Objective) packed in-house with ReproSil-Pur C18-AQ, 1.9 μm resin (Dr Maisch GmbH). Packed emitters were heated by means of a column oven (Sonation, Biberach, Germany) integrated into the nanoelectrospray ion source (Thermo Fisher Scientific, Waltham, MA, USA). An Active Background Ion Reduction Device (ABIRD, ESI Source Solutions, Woburn, MA, USA) was used to decrease air contaminants signal level. The Xcalibur software (Thermo Fisher Scientific, Waltham, MA, USA) was used for data acquisition.

An Orbitrap Fusion Lumos mass spectrometer (Thermo Fisher Scientific, Waltham, MA, USA) was used for the TMT-labelled proteome analysis. Peptides were loaded into a packed 50 cm fused silica emitter kept at 50°C, and eluted at a flow rate of 300 nl/min using different gradients optimised for three sets of fractions: 1–7, 8–15, and 16–21 as previously described (58). A full scan over mass range of 350–1400 m/z was acquired at 60,000 resolution at 200 m/z, with a target value of 500,000 ions for a maximum injection time of 20 ms. Higher energy collisional dissociation fragmentation was performed on the 15 most intense ions, for a maximum injection time of 100 ms, or a target value of 100,000 ions. Peptide fragments were analysed in the Orbitrap at 50,000 resolution.

AP-MS was carried out using an Orbitrap Q-Exactive HF mass spectrometer (Thermo Fisher Scientific, Waltham, MA, USA) using 20 cm fused silica emitter kept at 35°C to separate the peptides over a 60 minutes gradient at a flow rate of 300 nl/min. For the full scan a resolution of 60,000 at 200 m/z was used to scan the mass range from 350–1400 m/z. The top ten most intense ions in the full MS were isolated for fragmentation with a target of 50,000 ions, for a maximum of 75 ms, at a resolution of 15,000 at 200 m/z.

The MS Raw data were processed with MaxQuant software (59) version 1.6.3.3 (TMT proteome experiments) or 1.5.5.1 (AP-MS experiment) and searched with Andromeda search engine (60), querying SwissProt (61) Homo sapiens (30/04/2019; 42,438 entries). First and main searches were performed with precursor mass tolerances of 20 ppm and 4.5 ppm, respectively, and MS/MS tolerance of 20 ppm. Database was searched requiring specificity for trypsin cleavage and allowing maximum two missed cleavages. Methionine oxidation and N-terminal acetylation were specified as variable modifications, and Cysteine carbamidomethylation as fixed modification. The peptide, protein, and site false discovery rate (FDR) was set to 1 %.

For the TMT-proteome analysis MaxQuant was set to quantify on “Reporter ion MS2”, and TMT10plex was chose as Isobaric label. Interference between TMT channels were corrected by MaxQuant using the correction factors provided by the manufacturer. The “Filter by PIF” option was activated and a “Reporter ion tolerance” of 0.003 Da was used.

Proteins identified in the AP-MS experiment, were quantified according to the label-free quantification algorithm available in MaxQuant (62).

MaxQuant outputs were analysed with Perseus software version 1.6.2.3 (63). From the ProteinGroups.txt file, Reverse and Potential Contaminant flagged proteins were removed, as well as protein groups identified with no unique peptides. Only proteins quantified in three out of three replicate experiments, were included in the analysis. To determine significant change in protein abundance, a Student t-test with a 5% FDR (permutation-based) was applied using the corrected reporter ions Intensities for the TMT proteome analysis or label free quantitation intensities for the AP-MS experiment.

### Lipidomic analysis

Lipidomic analysis was performed according to (15). Single-phase lipid extraction was carried out using an extracting solution of methanol-butanol (1:1 ratio, BuMe), kept at 4°C and supplemented with an internal standard (Splash Lipidomix, Avanti Polar Lipids, Alabaster, AL, USA) further used as a quality control. Briefly, plates were washed twice with ice-cold PBS and incubated with BuMe on dry ice for 20 minutes. Extracted lipids were then transferred to 1.5 ml tubes and centrifuged for 10 minutes at 14,000 rpm (4°C). Supernatants were collected and stored at -80°C prior to LC-MS analysis. Protein pellets attached to the plate were left to dry overnight and quantified for subsequent normalisation. Lipid analyses were performed using an LC-MS system including an Ultimate 3000 HPLC (Thermo Fisher Scientific, Waltham, MA, USA) coupled to a Q-Exactive Orbitrap mass spectrometer (Thermo Fisher Scientific, Waltham, MA, USA). Polar and non-polar lipids were separated on an Acquity UPLC CSH C18 column (100 × 2.1 mm; 1.7 µm) (Waters corporation, Milford, MA, USA) maintained at 60°C. The mobile phases consisted of 60:40 ACN: H2O with 10 mM ammonium formate, 0.1% formic acid and 5 µM of phosphoric acid (A) and 90:10 IPA:ACN with 10 mM ammonium formate, 0.1% formic acid and 5 µM phosphoric acid (B). The gradient was as follows: 0-2 min 30% (B); 2-8 min 50% (B); 8-15 min 99% (B), 15-16 min 99% (B), 16-17 min 30% (B). Sample temperature was maintained at 6°C in the autosampler and 5 µL of sample were injected into the LC-MS instrument.

Thermo Q-Exactive Orbitrap MS instrument was operated in both positive and negative polarities, using the following parameters: mass range 240-1200 m/z (positive) and 240-1600 (negative), spray voltage 3.8 kV (ESI+) and 3 kV (ESI-), sheath gas (nitrogen) flow rate 60 units, auxiliary gas (nitrogen) flow rate 25 units, capillary temperature (320°C), full scan MS1 mass resolving power 70,000. Data dependent fragmentation (dd-MS/MS) parameters for each polarity as follows: TopN: 10, resolution 17,500 units, maximum injection time: 25 ms, automatic gain control target: 5e^5^ and normalised collision energy of 20 and 25 (arbitrary units) in positive polarity. TopN: 5, resolution 17,500 units, maximum injection time: 80 ms automatic gain control target: 5e^5^ and normalised collision energy of 20 and 30 (arbitrary units) in negative polarity. The instrument was externally calibrated to <1ppm using ESI positive and negative calibration solutions (Thermo Fisher Scientific, Waltham, MA, USA).

Feature detection and peak alignment from .Raw files were performed using Compound Discoverer 3.2 (Thermo Fisher Scientific, Waltham, MA, USA). Files were also converted to .mgf format using MSConvert software (ProteoWizard) and MS2 files were searched against the LipiDex_ULCFA database using LipiDex software (64).

Data pre-processing, filtering and basic statistics were performed using Perseus software version 1.6.14.0 after importing the final results table from LipiDex. For statistical purposes, we only kept lipid ions where the number of features was present in at least 60% of the samples. GraphPad Prism 8.4.2 was used to produce heatmaps.

### GC-MS-based determination of ^13^C-sterols and fatty acids

Fatty acids (FA) and sterol measurements were performed as previously described in and (65) respectively. Briefly, cells were incubated for 72 hours in glucose-and glutamine-free RPMI (Gibco, Thermo Fisher Scientific, Waltham, MA, USA) supplemented with 10% CSS, 2 mM [U13C]-glutamine and 10 mM [U13C]-glucose (Cambridge Isotopes Laboratories, UK).

FA were extracted in a methanol-chloroform-PBS buffer (750 µl 1:1 v/v PBS: methanol and 500 µl chloroform) supplemented with 50 µl of 1 mg ml^-1^ methanolic butylated hydroxytoluene (BHT, Sigma Aldrich, St Louis, MI, USA) and 20 µl of 0.05 mg ml^-1^ 17:0 PC (Avanti Polar Lipids, Albaster, AL, USA), used as an internal standard. Samples were centrifuged at 10,000 × g for 5 min before the lower chloroform layer was extracted and dried under N2. Samples were reconstituted in 90 µl chloroform and incubated with 10 µl MethPrepII (Thermo Fisher Scientific, Waltham, MA, USA) for 20 min at room temperature.

Sterols were extracted in a methanol extraction buffer (1:9 v/v water: methanol) supplemented with 20 μL of lathosterol (100 ng/μL) internal standard. Saponification was performed by heating samples for 60 min at 80°C with 75 μL of 10 mol/L NaOH to obtain the total cholesterol pool. Upon cooling to room temperature, 200 μL water and 500 μL n-hexane were sequentially added. Samples were then vortexed and the upper hexane layer was transferred to an autosampler vial. The n-hexane extraction was repeated and samples dried under N2. Samples were reconstituted in 50 μL dry pyridine and 50 μL N-Methyl-N-(trimethylsilyl) trifluoroacetamide (MSTFA, Sigma Aldrich, St Louis, MI, USA) silylation agent added. Samples were heated at 60°C for 60 minutes before cooling and immediate analysis by gas chromatography–mass spectroscopy (GC-MS).

Fatty Acid Methyl Esters (FAMEs) and sterols were analysed using an Agilent 7890B GC system coupled to a 7000 Triple Quadrupole GC-MS system, with a Phenomenex ZB-1701 column (30 mm × 0.25 mm × 0.25 μm). For FAMEs, an initial temperature of 45°C was set to increase at 9°C/min, held for 5 min, then 240°C min^-1^, held for 11.5 min, before reaching a final temperature of 280°C min^-1^, held for 2 min. For cholesterol, the initial temperature was set at 200°C and increased at 20°C/minute up to 280°C, and held for 9 minutes. The instrument was operated in pulsed splitless mode in the electron impact mode, 50eV, and mass ions were integrated for quantification using known standards to generate a standard curve. Palmitic, stearic and oleic acid peak areas were extracted using mass-to-charge ratios (m/z) 270, 298 and 296 respectively. Cholesterol peak areas were extracted from m/z 458. Mass Hunter B.06.00 software (Agilent) was used to quantify isotopomer peak areas before natural abundance isotope correction was performed using an in-house algorithm.

Cholesterol was analyzed using an Agilent 7890B GC system coupled to an Agilent 7000 Triple Quadrupole GC-MS system, which was operating in a single quadrupole mode, with a Phenomenex ZB-1701 column (30 mm × 0.25 mm × 0.25 μm). An initial temperature of 200°C was set to increase at 20°C/minute up to 280°C, and held for 9 minutes. The instrument was operated in splitless mode in the electron impact mode, 70eV, for quantification and 50eV for labeling experiments. Cholesterol was quantified and isotope labeling pattern analyzed using Mass Hunter B.06.00 software (Agilent). Cholesterol and lathosterol internal standard peak areas were extracted from mass-to-charge ratio (m/z) 458 for both. Cholesterol was normalized to the internal standard, and a standard curve was used to quantify mg cholesterol per sample.

### Bioinformatics analysis

Gene expression data were downloaded from TCGA and the GEO website. TCGA RNASeqV2 data were shift log transformed; GSE35988 and GSE21034 data were log transformed, using mean of probes per gene. Expression values were grouped according to sample type and group distributions were plotted using the matlab routine boxplot with the bar indicating the median, the box spanning from the 25th to the 75th percentiles, and whiskers spanning 2.7\sigma. Outliers beyond that span are indicated in red. Significant difference between the groups was measured using overall ANOVA.

For survival analysis (GSE21034), gene expression data from human samples (excluding cell lines) was normalised as before, mean-centered, and clustered into three groups using kmeans. Kaplan-Meier survival curves for the groups were plotted using the matlab routine kmplot.

### Human prostate and breast cancer orthografts

*In vivo* orthograft experiments were performed in accordance with the ARRIVE guidelines (66), and by a local ethics committee (University of Glasgow) under the Project Licences P5EE22AEE and 70/8645 in full compliance with the UK Home Office regulations (UK Animals (Scientific Procedures) Act 1986) and EU directive. For PCa cells, 10×10^6^ cells/mouse were suspended in a 1:1 solution of serum-free medium and growth factor reduced Matrigel (Corning, NY, USA). 50 µl of cell suspension were injected orthotopically into the anterior prostate lobe of 10-weeks old CD1-*nude* male mice (Charles River Laboratories, UK). Orchidectomy was either performed at the time of injection (22rv1) or 3 weeks after cell implantation (CWR22res). Tumours were allowed to develop for 8 weeks. Tumour volume was measured weekly using a Vevo3100 ultrasound imaging system (Fujifilm Visualsonics, The Netherlands). For BCa cells, 1×10^6^ cells/mouse were suspended in 50 µl of a 1:1 solution of serum-free medium and growth factor reduced Matrigel (Corning, NY, USA) and injected into the mammary fat pad of 8-weeks old CD1-*nude* female mice (Charles River Laboratories, UK). Tumours were allowed to grow for 35 days. Once the tumour was palpable, tumour volume was calliper measured twice a week with volume calculated using the formula (width² × length)/2. At the end of the experiment, tumours were collected, cut into pieces and either fixed in 10% formalin for histological procedures or snap-frozen in liquid nitrogen.

### Raman Spectroscopy analysis

Raman spectra were acquired on a Renishaw inVia Raman microscope equipped with a 532 nm Nd:YAG laser giving a maximum power of 500 mW, 1800 l mm^-1^ grating, and a Nikon NIR Apo 60×/1.0 N.A. water dipping objective. Prior to Raman measurements, tissue sections were dewaxed in xylene (2 × 15 min), 100% ethanol (5 min), 95% EtOH (5 min) and 90% EtOH (5 min). A water dipping objective was used to map the tissue sections with a step size of 100 µm in x and y, with 1 s acquisition time, 100% laser power and a spectral center of 3000 cm^−1^.

For initial pre-processing steps Renishaw Wire 4.1 software was used. The inbuilt software functions were used to remove cosmic rays followed by baseline subtraction. Baseline was subtracted using the baseline subtraction intelligent fitting function (with an 11^th^ order polynomial fitting and noise tolerance set to 1.50, applied to the whole spectral dataset). Further data analysis steps were then performed using custom MATLAB® scripts. Outlier spectra which gave high intensity due to saturation or fluorescence were removed using a threshold function. Spectra were cut to the region of interest between 2800 cm^−1^ and 3020 cm^−1^. Spectra from tissue regions were then extracted (based on total spectral intensity) and all extracted spectra were min-max scaled for comparison between conditions. For each map the resultant spectral data set went through further quality control steps which involved firstly, excluding spectra with a total spectral intensity less than 60000, and then removing spectra out with one standard deviation of the mean. Spectra were then scaled to the peak at 2933 cm^−1^, and spectral data sets for all CTL and THEM6 KO samples were combined. The average for each condition was plotted for comparison. The ratio of the intensities of the peaks at 2845 cm^−1^ and 2935 cm^−1^ as well as the ratio of the intensities of the peaks at 2880 cm^−1^ and 2935 cm^−1^ were determined for each spectral data point. GraphPad Prism 8.4.2 was used to produce graphs and perform statistical analysis of the data.

### Patient material and immunohistochemistry

This study was approved by the West of Scotland Research Ethics Committee (05/S0704/94) and the Chair of the University of British Columbia Clinical Research Ethics Board (UBC CREB number: H09-01628). All patients involved in this study provided written informed consent and all experiments conformed to the principles set out in the WMA Declaration of Helsinki and the Department of Health and Human Services Belmont Report.

Immunohistochemical (IHC) staining for THEM6 was performed on 4 µm formalin-fixed paraffin-embedded sections which had previously been incubated at 60°C for 2 hours. The IHC staining was performed on an Agilent Autostainer link 48 (Agilent, Santa Clara, CA, USA) or on a Ventana DISCOVERY Ultra (Ventana Medical Systems, Tucson, Arizona, USA). Briefly, formalin-fixed paraffin-embedded (FFPE) TMA sections were deparaffinised before undergoing antigen retrieval. Sections were then successively incubated with anti-THEM6 antibody (rabbit, 1:5000, ab121743, Abcam, Cambridge, UK) and respective secondary antibody before being detected using Liquid DAB (Agilent, Santa Clara, CA, USA) or UltraMap DAB anti-Rb Detection Kit (Ventana Medical Systems, Tucson, Arizona, USA). Stained slides were scanned with Leica Aperio AT2 (Leica Microsystems, Concord, Ontario, Canada). The area of interest in the tumour images were delineated by pathologist. Positively stained cells were quantified with Aperio ImageScope (Leica Biosystems, Buffalo Grove, Illinois, USA).

### Electron microscopy

CTL and THEM6 KO 22rv1 cells were fixed for 2 h at 4°C in a solution composed of 2.5% glutaraldehyde in 0.1 M Sorensen’s buffer (0.2 M NaH2PO4, 0.2 M Na2HPO4, pH 7.4). After several washes in the same buffer, the samples were post-fixed for 60 min with 2% osmium tetroxide, washed in deionized water, dehydrated through graded ethanol (70%, 95%, and 100%), and embedded in epon for 48 h at 60°C. Ultrathin sections (700-A thick) were obtained by means of an ultramicrotome (Reichert Ultracut E) equipped with a diamond knife. The ultrathin sections were mounted on palladium/copper grids coated with collodion and contrasted with uranyl acetate and lead citrate for 5 min each before being examined under a Jeol JEM1400 transmission electron microscope at 80 kV.

### siRNA transfection

70%-confluent cells were transfected using Lipofectamine RNAimax (siRNA) or Lipofectamine 2000 (plasmid) (Invitrogen, Thermo Fisher Scientific, Waltham, MA, USA) according to the manufacturer’s protocol. ON-TARGETplus smartpool siRNAs against *hTHEM6* (L-020791-01-0005), *mThem6* (L-057277-01-0005) as well as non-targeting siRNA (D-001810-01-20) were purchased from Dharmacon (Dharmacon, Horizon inspired cell solutions, Cambridge, UK). *THEM6* human tagged ORF clone overexpressing plasmid and respective control (RC201709 and PS100001, Origene, Rockville, MD, USA) were purchased from Origene. Protein and RNA extractions were performed 48 hours after transfection.

### qPCR analysis

Total RNA was extracted from 80%-confluent cells using the RNeasy Mini Kit (Qiagen, Hilden, Germany) with on-column DNase digestion (RNase-Free DNase Set, Qiagen, Hilden, Germany). 4 ug of RNA were then used for cDNA preparation using the High-Capacity cDNA Reverse Transcription Kit (Thermo Fisher Scientific, Waltham, MA, USA). Real-time PCR was performed using TaqMan Universal Master Mix (Thermo Fisher Scientific, Waltham, MA, USA) with primer-appropriate Universal ProbeLibrary probes (Roche, Basel, Switzerland) and the ABI 7500 FAST qPCR system (Thermo Fisher Scientific, Waltham, MA, USA). *CASC3* gene was used as a normalising control. Gene expression is shown relative to control cell levels. Primers (Thermo Fisher Scientific, Waltham, MA, USA) used in this study are provided in Supplementary Table 5.

### Immunoblotting

20 µg of proteins extracted in SDS buffer (1% SDS supplemented with protease and phosphatase inhibitors) were loaded on to a 4-12% gradient SDS-PAGE gel (Invitrogen, Thermo Fisher Scientific, Waltham, MA, USA) and transferred to a PVDF membrane (GE Healthcare, Chicago, IL, USA). Membrane was sequentially probed overnight with primary antibodies (see Supplementary Table 6) diluted in 5% BSA-TBST, and respective HRP-conjugated secondary antibodies diluted in 5% milk-TBST. Antigen revelation was performed using the ECL kit (GE Healthcare, Chicago, IL, USA) and images were acquired on a MyECL machine (Thermo Fisher Scientific, Waltham, MA, USA).

### Immunofluorescence

Cells seeded on glass coverslips were fixed in ice-cold Methanol/Acetone buffer, washed, blocked in 5% BSA and probed with primary antibodies overnight (anti-THEM6, ab121743, anti-CALR, ab22683, Abcam, Cambridge, UK; anti-mitochondria, MAB1273, Merck Millipore, Burlington, MA, USA). The next day, coverslips were washed and incubated with fluorophore-coupled secondary antibodies (Abcam, Cambridge, UK). Coverslips were mounted using Diamond Prolong with DAPI (Thermo Fisher Scientific, Waltham, MA, USA). Pictures were taken on a Nikon A1R confocal microscope (Nikon Instruments Europe B.V., Amsterdam, The Netherlands).

### Statistical analysis

Statistical analyses were performed using GraphPad PRISM software v8.4.2 (GraphPad Software Inc, San Diego, CA, USA).

### Data reproducibility

Figure 1: Panel **A**: n = 3 tumours per group. Panel **B**: n = 1 gel loaded with three prostate orthografts per condition. Panels **C, K:** representative image from 3 independent biological experiments. Panel **D, E**: n = 132; 90; 66; 137; 44; 30 for Untreated; NHT 1-3; NHT4-6; NHT>7; CRPC; NEPC respectively. Panel **F**: n = 99 for Gleason score ≤ 7 and n = 63 for Gleason score > 7. Panel **G**: n = 151 for non-metastatic patients and n = 76 for metastatic patients. Panels **H, I**: n = 35 for THEM6 low and n = 34 for THEM6 high. Panel **J**: n = 269 for Ki67 low and n = 270 for Ki67 high. Panel **L**: n = 3 independent biological experiments. Panels **M, N**: n = 5 mice per group.

Figure 2: Panels **A, B, C**: n = 3 independent biological experiments. Panel **D**: n = 4 mice per group. Panel **E, F**: n = 6581 (CTL); 12047 (KO1); 6493 (KO2) peak intensities that were extracted from 4 mice per group.

Figure 3: Panel **A** (left): n = 46 (CTL); 47 (KO1); 49 (KO2) cells measured. Panel **A** (right): n = 31 (CTL); 43 (KO1); 50 (KO2) cells measured. Panels **E, G, K**: representative image from 3 independent biological experiments. Panel **J**: representative image from 2 independent biological experiments.

Figure 4: Panel **B**: n = 3 independent biological experiments. Panels **C, D, G**: n = 3 independent wells from the same cell culture. Panel **F**: n = 225 for THEM6 low and n = 121 for THEM6 high.

Figure 5: Panels **C, F, G, I, J**: representative image from 3 independent biological experiments. Panel **D**: n = 3 independent biological experiments. Panel **H**: n = 225 for THEM6 low and n = 121 for THEM6 high.

Figure 6: Panel **B**: n = 489 patients. Panel **C**: n = 1 gel loaded with three prostate orthografts per condition. Panel **D**: representative image from 2 independent biological experiments. Panel **E, G**: n = 3 independent biological experiments. Panels **F, L, N**: representative image from 3 independent biological experiments. Panel **H**: n = 307 for LN negative patients and n = 234 for LN positive patients. Panel **I**: n = 93; 238; 220 for patients with low, medium and high tumour invasiveness respectively. Panel **J**: n = 362 for Ki67 low and n = 189 for Ki67 high. Panel **K**: n = 319 for THEM6 low and n = 491 for THEM6 high. Panel **M**: n = 101 for THEM6 low and n = 22 for THEM6 high. Panel **O**: n = 48 (CTL); 49 (KO1); 43 (KO2) cells measured. Panel **P**: n = 7 (CTL) and 8 (KO) mice per group.

## DATA AVAILABILITY

For proteomics, the raw files and the MaxQuant search results files have been deposited as partial submission to the ProteomeXchange Consortium via the PRIDE partner repository (67). For lipidomics, the raw LC-MS/MS files and the processed data have been deposited on the Metabolomics Workbench repository.

The following databases were used in this study: The Cancer Genome Atlas (TCGA - https://tcga-data.nci.nih.gov/tcga/); GSE35988 (https://www.ncbi.nlm.nih.gov/geo/query/acc.cgi?acc=GSE35988);

GSE21034 (https://www.ncbi.nlm.nih.gov/geo/query/acc.cgi?acc=GSE21034); STRING v11.0 (https://string-db.org/cgi/input.pl).

## ACKNOWLEDGEMENTS

We would like to thank the Core Services and Advanced Technologies at the Cancer Research UK Beatson Institute, with particular thanks to the Metabolomics and Proteomics Units, the Biological Services and Histology facilities of the Beatson Institute CRUK, as well as the PRIDE team. This work was supported by Cancer Research UK Beatson Institute core funding (C596/A17196) and CRUK core group awarded to HYL (A15151), to KB (A29799) and to SZ (A29800). DJM was supported by a CRUK EDx grant (C48702/A27603). MS is a Medical Research Council Clinical Research Fellow (MR/L017997/1). CN is the recipient of CRUK Clinical Research Fellowship (grant 300444-01). DG and KF acknowledge support from the EPSRC grant EP/L014165/1 that supported LJ.

## AUTHOR CONTRIBUTIONS

AB and HYL designed the study. AB, CP, EM, GRB, LMM, LEJ, GHM, SL, MT, SHYK, CAF, LKR, DJM, SM, CNi, MJS performed the experiments. AB, GRB, LMM, LEJ, GHM, SL, CN, MT, SHYK, PR, EMa, GMM, JE, DS analysed the data. AB, RP, MT, JJK, DG, KF, LF, MEG, EA, JE, HY, KB, PC, DJMu, SZ and HYL interpreted and discussed the data. AB and HYL wrote the manuscript. All authors critically reviewed the manuscript.

## SUPPLEMENTARY FIGURE LEGENDS

**Supplementary Figure 1:**
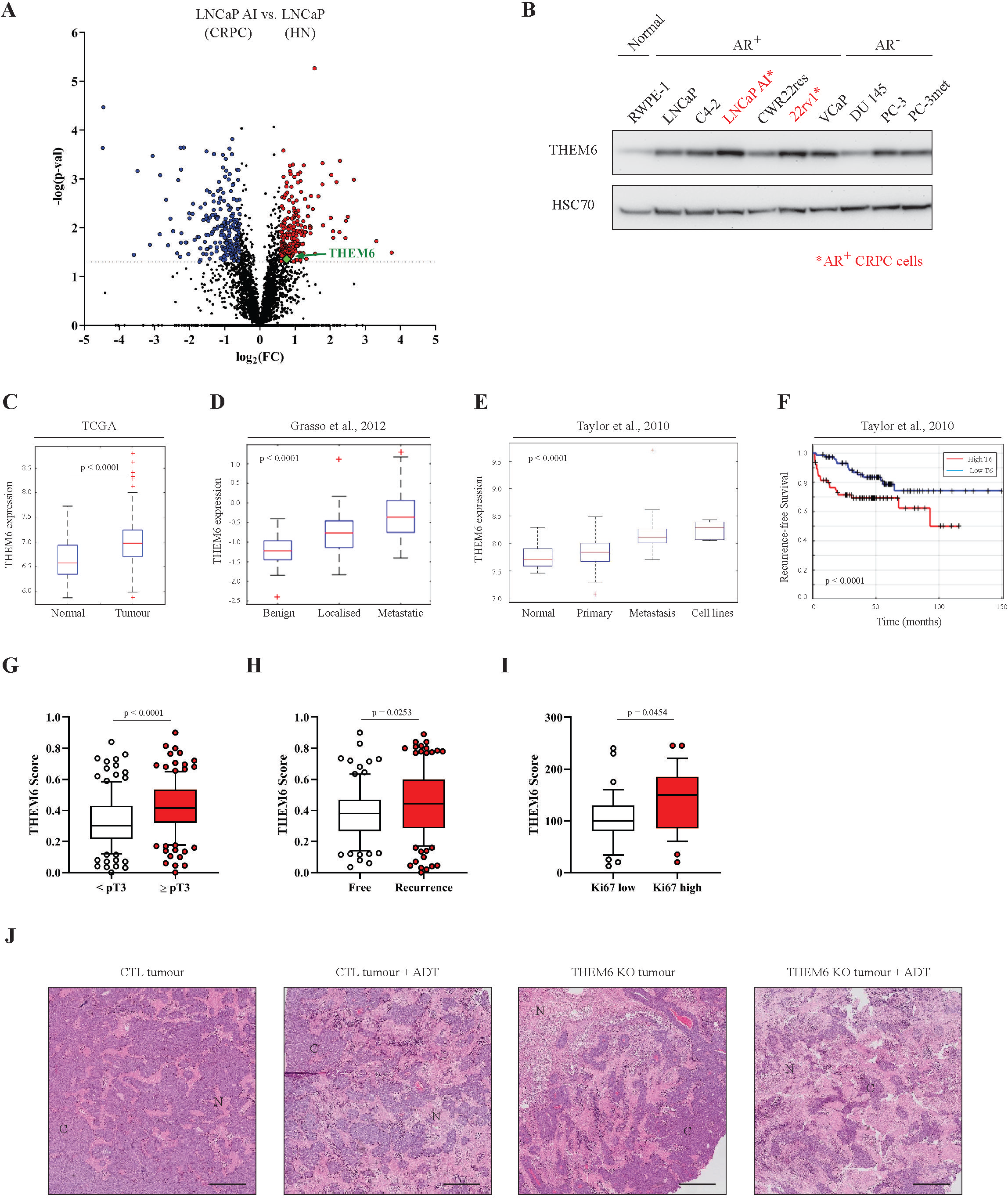
THEM6 is a clinically relevant protein overexpressed in CRPC. **A**, Volcano plot of the differentially modulated proteins in LNCaP AI (CRPC) versus LNCaP (HN) tumours. Red and blue dots represent the proteins that are significantly up- and down-regulated, respectively (p-value ≤ 0.05, FC = 1.5). **B**, Western blot analysis of THEM6 expression in PCa cells. HSC70 was used as a sample loading control. **C**, Gene expression analysis of THEM6/c8orf55 in normal and tumoural prostate tissues according to the PRAD TCGA dataset (n=489). **D**, Gene expression analysis of THEM6/c8orf55 in benign, localised and metastatic tumoural prostate tissues according to the GSE35988 dataset (n=122). **E**, Gene expression analysis of THEM6/c8orf55 in normal prostate tissues, primary tumours, distant metastases and PCa cell lines according to the GSE21034 dataset (n=179). **F**, Kaplan-Meier recurrence-free survival analysis of PCa patients stratified according to median THEM6 expression using the GSE21034 dataset. **G-I**, Quantification of THEM6 expression in PCa tissue samples according to T-stage (**G**), cancer recurrence (**H**) and Ki67 score (**I**). **J,** Representative pictures of hematoxylin/eosin staining on orthografts from CWRres CTL and THEM6 KO tumours. C = cancer cells; N = necrotic area. Scale bar represents 1000 µm. Panels **C, D, E**: Center line corresponds to median of data, top and bottom of box correspond to 75th and 25th percentile, respectively. Whiskers extend to adjacent values (minimum and maximum data points not considered outliers). Panels **G, H, I**: Center line corresponds to median of data, top and bottom of box correspond to 90th and 10th percentile, respectively. Whiskers extend to adjacent values (minimum and maximum data points not considered outliers). <u>Statistical analysis</u> : **C, D, E**: pairwise ANOVA. **F**: logrank test. **G, H, I**: two-tailed Mann-Whitney U test. <u>Data reproducibility</u>: **A**: n = 3 tumours per group. **B**: representative image from 3 independent biological experiments. **G**: n = 129 for T stage score < 3 and n = 135 for T stage score ≥ 3. **H**: n = 91 for disease-free patients and n = 136 for recurred patients. **I**: n = 35 for Ki67 low and n = 29 for Ki67 high. **J**: representative image from 3 tumours per group.

**Supplementary Figure 2:**
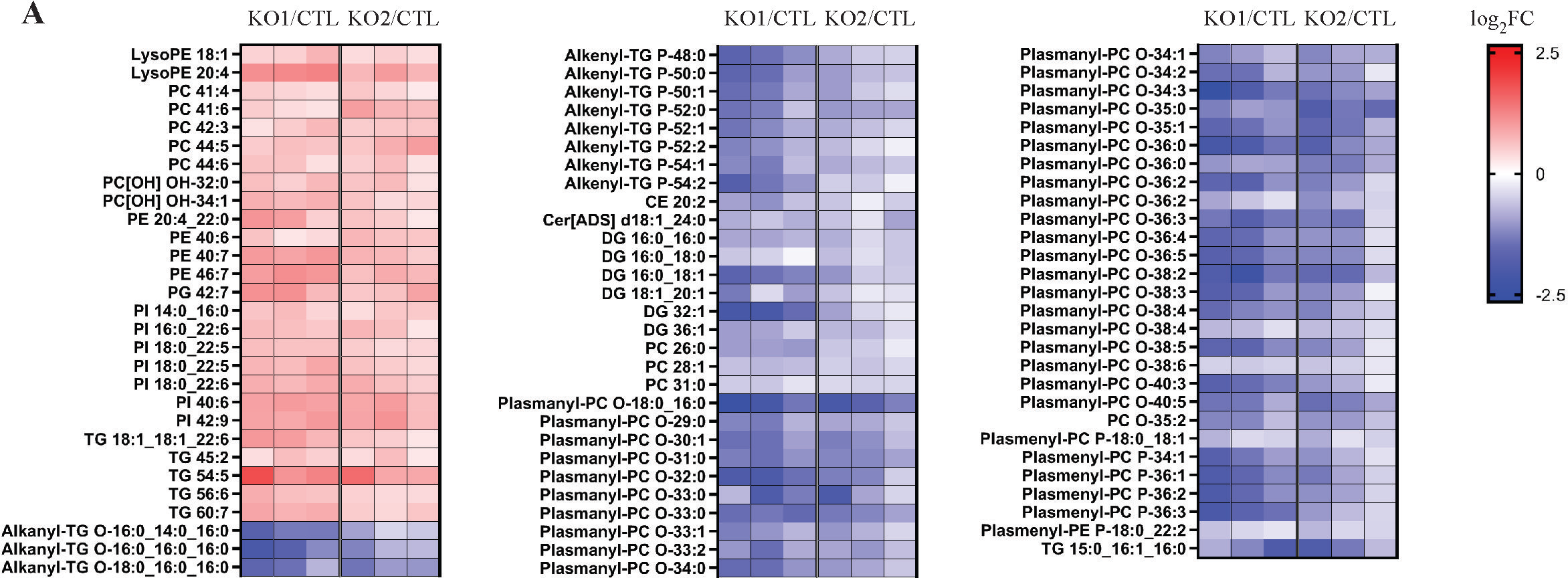
Loss of THEM6 alters lipid homeostasis. **A,** Heatmap illustrating the steady-state levels of significantly regulated lipids in THEM6 KO LNCaP AI cells when compared to CTL (p ≤ 0.05, FC = 1.3). Values are expressed as log2(FC). <u>Data reproducibility</u>: **A**: n = 3 independent biological experiments.

**Supplementary Figure 3:**
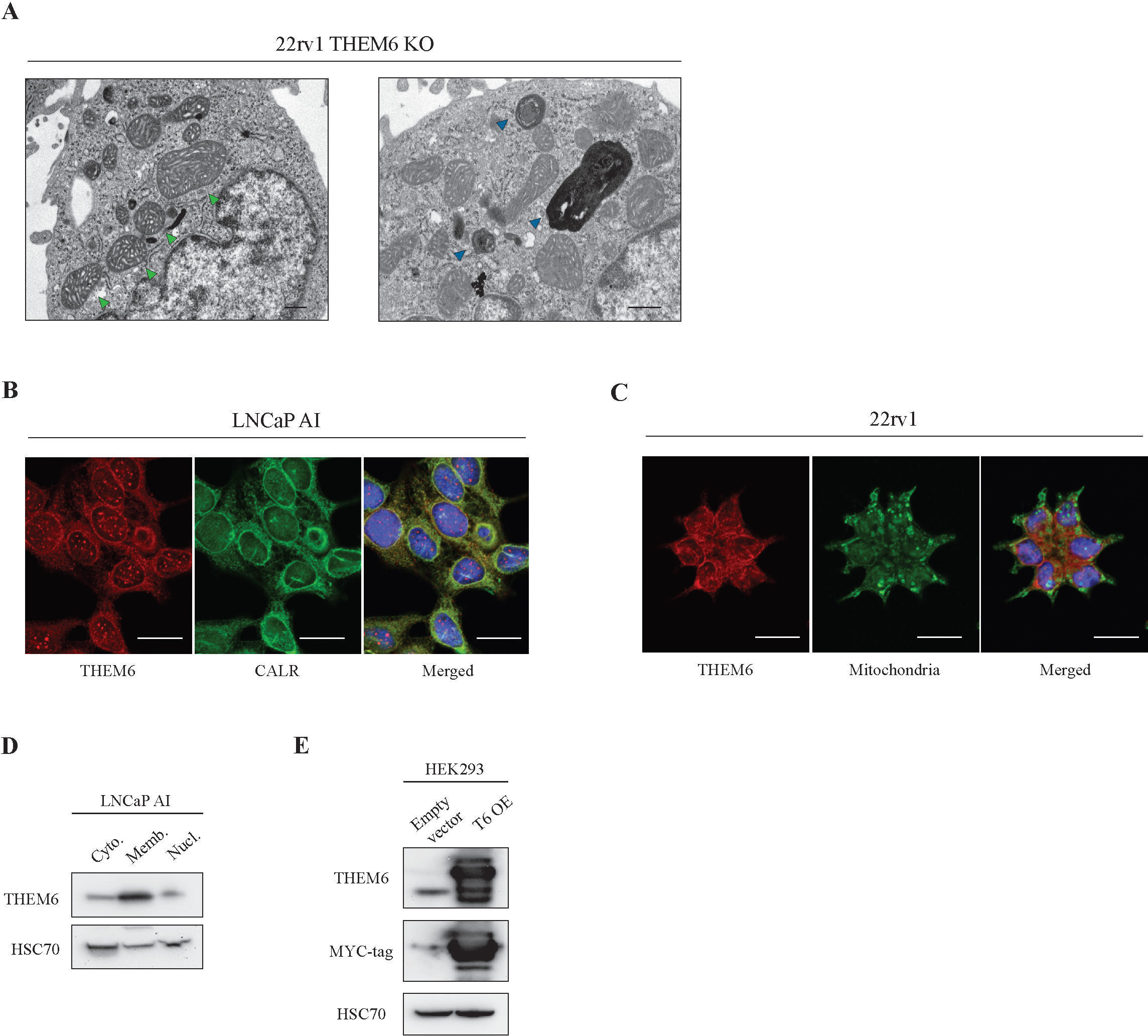
THEM6 is an ER membrane protein. **A,** Representative electron microscopy pictures of THEM6 KO 22rv1 cells. Green and blue arrows point towards enlarged mitochondria and abnormal lysosomal structure, respectively. Scale bar represents 500 nm. **B,** Immunofluorescence showing co-localisation of THEM6 and the ER marker calreticulin in LNCaP AI cells. Scale bar represents 20 µm. **C,** Immunofluorescence showing distinct localisations for THEM6 and mitochondria in 22rv1 cells. Scale bar represents 20 µm. **D**, Western blot analysis of THEM6 expression in cytoplasmic (cyto.), membrane/organelle (memb.) and nuclear fractions (nucl.) of LNCaP AI cells. HSC70 was used as cytoplasmic-enriched marker. **E**, Western blot analysis of THEM6 and MYC-tag expression in HEK293 cells overexpressing a MYC-tagged version of THEM6 (T6 OE). HSC70 was used as a sample loading control. <u>Data reproducibility</u>: **B, C, D**: representative image from 3 independent biological experiments. **E**: representative image from two independent biological experiments.

**Supplementary Figure 4:**
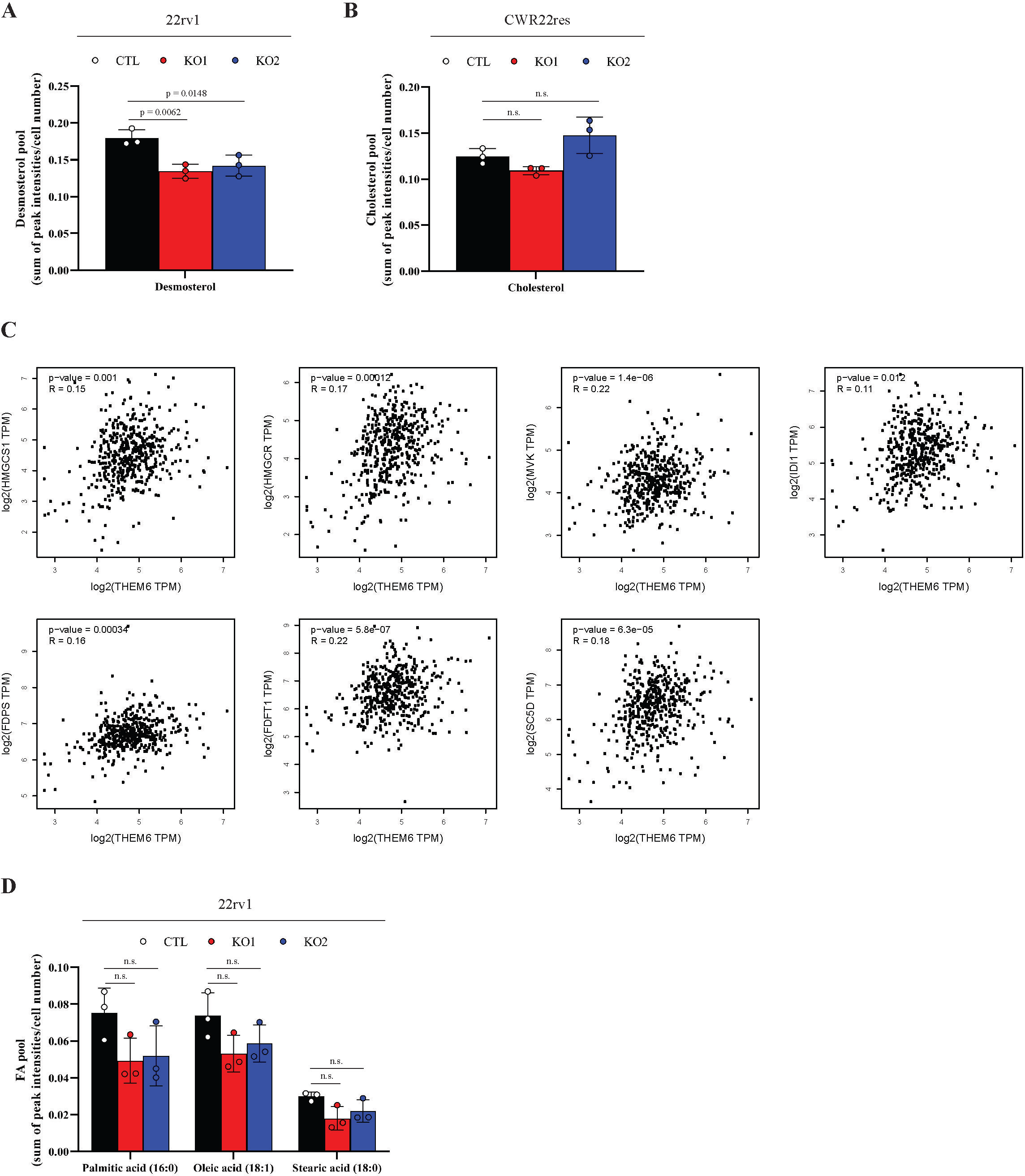
THEM6 expression weakly correlates with enzymes involved in the early steps of sterol biosynthesis. **A,** Total pool of desmosterol in CTL and THEM6 KO 22rv1 cells. Data extracted from Fig. 4c. **B,** Total pool of cholesterol in CTL and THEM6 KO CWR22res cells. Data extracted from Fig. 4d. **C,** Pearson’s correlation analysis of *HMGCS1, HMGCR, MVK, IDI1, FDPS, FDFT1, SC5D* with *THEM6* using the PRAD TCGA dataset. Results were obtained using the GEPIA website http://gepia.cancer-pku.cn/. **D,** Total pool of palmitic, oleic and stearic acid in CTL and THEM6 KO 22rv1 cells. Data extracted from Fig. 4g. Panels **A, B, D**: Data are presented as mean values +/- SD. <u>Statistical analysis</u> : **A, B, D**: 1-way ANOVA with a Dunnett’s multiple comparisons test. <u>Data reproducibility</u>: **A, B, D**: n = 3 independent wells from the same cell culture.

**Supplementary Figure 5:**
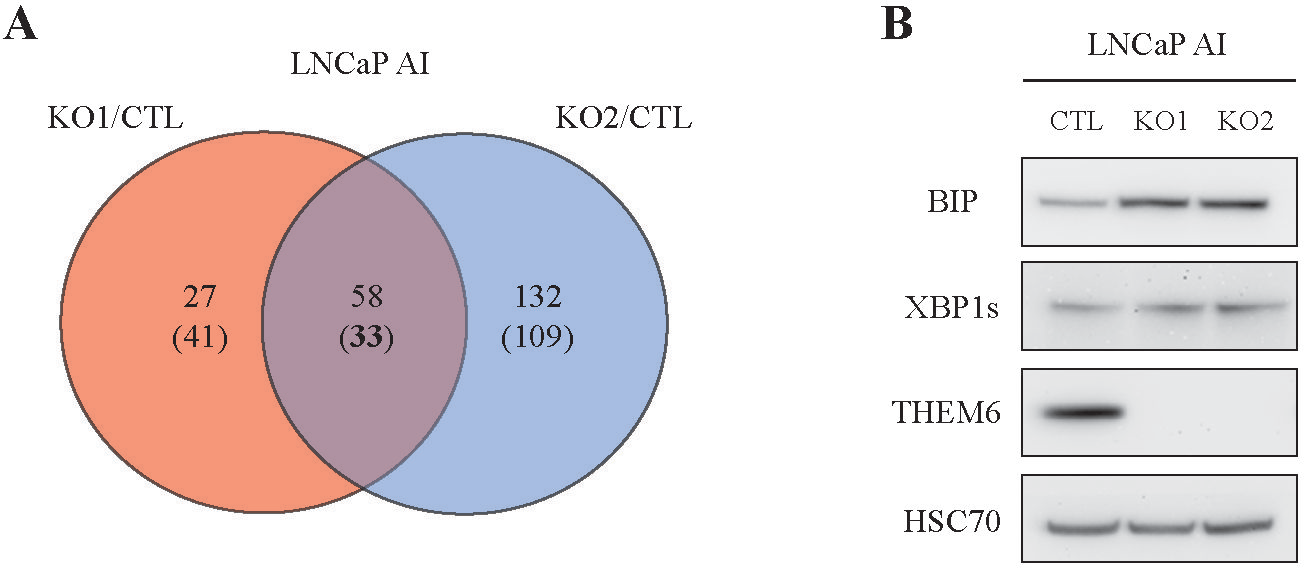
Proteomic analysis of THEM6 KO LNCaP AI cells. **A,** Venn diagram highlighting commonly modulated proteins (p-value ≤ 0.05, FC = 1.3) in THEM6 KO LNCaP AI cells (2 clones) when compared to CTL. Up-regulated proteins are on top; Down-regulated proteins are into brackets. **B**, Western blot analysis of BIP, XBP1s and THEM6 expression in CTL and THEM6 KO LNCaP AI cells. HSC70 was used as a sample loading control. <u>Data reproducibility</u>: **B**: representative image from 3 independent biological experiments.

**Supplementary Figure 6:**
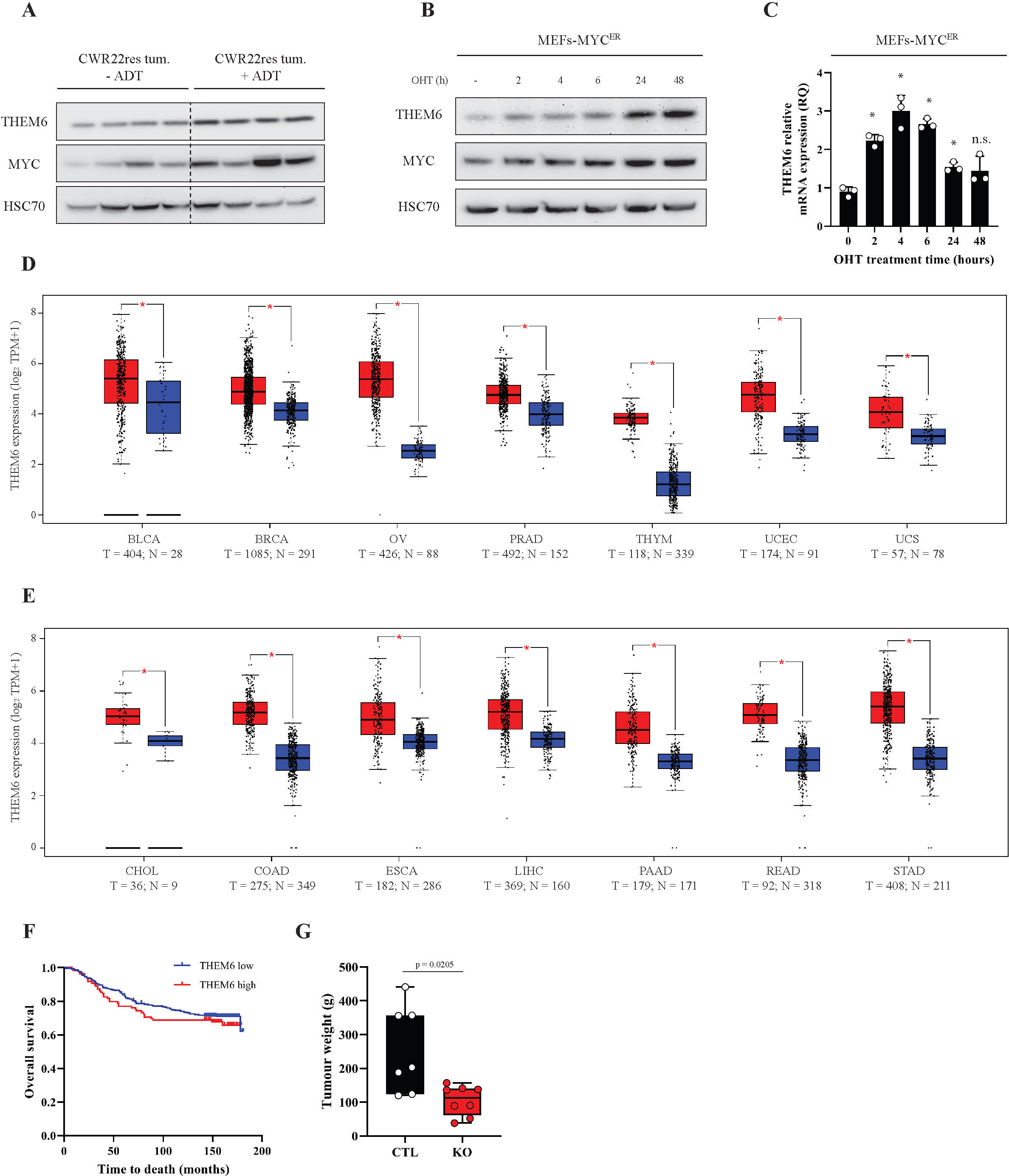
THEM6 is overexpressed in multiple cancer types. **A**, Western blot analysis of MYC and THEM6 expression in HN (-ADT) and CRPC (+ADT) CWR22res prostate orthografts. **B**, Western blot analysis of MYC and THEM6 expression in MEFs-MYC^ER^ along time. **C,** RT-qPCR analysis of *Them6* expression in MEFs-MYC^ER^ along time. *CASC3* was used as a normalising control. **D-E**, Gene expression analysis of THEM6/c8orf55 in normal and tumoural tissues from different origins, according to their respective TCGA datasets. **F**, Kaplan-Meier overall survival analysis of BCa patients stratified according to upper quartile THEM6 expression. **G**, Tumour weight of CTL and THEM6 KO MDA-MB-231-derived orthografts taken at endpoint. Panels **A, B**: HSC70 was used as a sample loading control. Panels **C, G**: Data are presented as mean values +/- SD. <u>Statistical analysis</u> : **C**: 1-way ANOVA with a Dunnett’s multiple comparisons test. **F**: logrank test. **G**: two-tailed Mann-Whitney U test. <u>Data reproducibility</u>: **A**: n = 1 gel loaded with four prostate orthografts per condition. **B**: representative image from 2 independent biological experiments. **C**: n = 3 technical replicates. **F**: n = 419 for THEM6 low and n = 132 for THEM6 high. **G**: n = 7 (CTL) and 8 (KO) mice per group.

## SUPPLEMENTARY TABLES

Supplementary Table 1: Proteins significantly modulated in THEM6 KO 22rv1 cells (2 clones).

Supplementary Table 2: Potential THEM6 interactors in MYC-OE HEK293 cells.

Supplementary Table 3: Proteins significantly modulated in high THEM6 tumours according to the PRAD TCGA dataset.

Supplementary Table 4: Proteins significantly modulated in THEM6 KO LNCaP AI cells (2 clones).

Supplementary Table 5: List of primers used in this study.

Supplementary Table 6: List of antibodies used in this study.

