## Supplementary Table 5 for "THEM6-mediated lipid remodelling sustains stress resistance in cancer"

**Supplementary Table 5:** List of primers used in this study.

| **Target Gene** | **Forward** | **Reverse** |
| --- | --- | --- |
| *hTHEM6* | ggagacaccaggctactaggac | tttccccagctgtaaggtga |
| *mTHEM6* | caagcaggccagagtagtca | cccactgtccctgagtaagc |
| *MVD* | gaccagggaaggggtcac | gcacttggtggtttcctga |
| *FDPS* | gagcggattctgcttttagg | gaagacccccacagatctca |
| *DHCR7* | aaaggggctttcatgtcgtt | cagactccaggcagagcac |
