## Supplementary Table 6 for "THEM6-mediated lipid remodelling sustains stress resistance in cancer"

| **Target** | **Species** | **Company** | **Reference** |
| --- | --- | --- | --- |
| THEM6 | Rabbit | abcam | ab121743 |
| VCL | Rabbit | Cell Signaling Technology | #13901 |
| HSC70 | Mouse | Santa Cruz | sc-7298 |
| CALR | Mouse | abcam | ab22683 |
| Mitochondria | Mouse | Millipore | MAB1273 |
| SP1 | Rabbit | Cell Signaling Technology | #9389 |
| AMFR | Rabbit | Cell Signaling Technology | #9590 |
| SEC61b | Rabbit | Cell Signaling Technology | #14648 |
| XPO1 | Rabbit | Cell Signaling Technology | #42649 |
| MYC-tag | Mouse | Cell Signaling Technology | #2276 |
| BiP | Rabbit | Cell Signaling Technology | #3177 |
| XBP1s | Rabbit | Cell Signaling Technology | #40435 |
| CHOP | Mouse | Cell Signaling Technology | #2895 |
| ATF4 | Rabbit | Cell Signaling Technology | #11815 |
| CALX | Rabbit | Cell Signaling Technology | #2679 |
| c-MYC | Rabbit | abcam | ab32072 |

**Supplementary Table 6:** List of antibodies used in this study.
